# Biased orientation representations can be explained by experience with non-uniform training set statistics

**DOI:** 10.1101/2020.07.17.209536

**Authors:** Margaret Henderson, John Serences

## Abstract

Visual acuity is better for vertical and horizontal compared to other orientations. This cross-species phenomenon is often explained by “efficient coding”, whereby more neurons show sharper tuning for the orientations most common in natural vision. However, it is unclear if experience alone can account for such biases. Here, we measured orientation representations in a convolutional neural network, VGG-16, trained on modified versions of ImageNet (rotated by 0°, 22.5°, or 45° counter-clockwise of upright). Discriminability for each model was highest near the orientations that were most common in the network’s training set. Furthermore, there was an over-representation of narrowly tuned units selective for the most common orientations. These effects emerged in middle layers and increased with depth in the network. Biases emerged early in training, consistent with the possibility that non-uniform representations may play a functional role in the network’s task performance. Together, our results suggest that biased orientation representations can emerge through experience with a non-uniform distribution of orientations, supporting the efficient coding hypothesis.

## Introduction

Contrary to common intuition, visual perception is not perfectly uniform across orientation space. One example of this principle is the “oblique effect”, which has been demonstrated in humans and a wide range of animal species, including cats, octopi and goldfish, among others. This effect describes the finding that observers’ ability to discriminate small changes in orientation, as well as other forms of acuity, tend to be worst for stimuli that have edges oriented diagonally (oblique orientations) and better for stimuli with edges oriented vertically or horizontally (cardinal orientations) (Appelle, 1972; Bauer, Owens, Thomas, & Held, 1979). In visual cortex, this finding has been linked to a larger number of orientation tuned neurons with a preference for cardinal orientations, as has been shown in cats (Li, Peterson, & Freeman, 2003), and macaques (Mansfield, 1974; Shen et al., 2014), among other species. Some evidence also suggests that cardinally-tuned neurons may have narrower tuning than other orientations, which may also contribute to higher acuity (Kreile, Bonhoeffer, & Hübener, 2011; Li et al., 2003).

One compelling explanation for the origin of the oblique effect is the efficient coding hypothesis, which suggests that because the brain operates with limited resources, coding resources should be preferentially allocated to stimuli that are highly probable during natural vision (Barlow, 1961; Girshick, Landy, & Simoncelli, 2011). On this view, biased orientation perception may reflect an adaptation to the statistics of natural images, in which vertical and horizontal orientations are most common (Coppola, Purves, McCoy, & Purves, 1998; Girshick et al., 2011). Support for an experience-driven account of the oblique effect includes evidence that in primates, the over-representation of cardinal orientations in visual cortex increases with age (Shen et al., 2014). Additionally, exposing developing kittens or mice to an environment with contours of only one orientation can induce changes in the distribution of cortical orientation tuning, suggesting some degree of plasticity (Blakemore & Cooper, 1970; Hirsch & Spinelli, 1970; Kreile et al., 2011; Leventhal & Hirsch, 1975). Finally, past computational work demonstrates that various models of optimal information coding, along with measurements of environmental statistics, can be used to predict both the neural and behavioral correlates of the oblique effect (Ganguli & Simoncelli, 2011; Girshick et al., 2011; Wainwright, 1999).

In addition, innate factors may also contribute to the efficient coding of cardinal orientation. For instance, while it is possible to significantly modify the distribution of orientation tuning preferences in visual cortex through experience, exposing an animal to only diagonal lines during development does not entirely obliterate tuning for cardinal orientations (Kreile et al., 2011; Leventhal & Hirsch, 1975). Similarly, rearing animals in complete darkness can result in a more extreme over-representation of cardinal-tuned units (Leventhal & Hirsch, 1980). In both mice and ferrets, it has been suggested that innate factors result in a strong oblique effect early in development, while visual experience tends to make orientation tuning more uniform over time (Coppola & White, 2004; Hoy & Niell, 2015). These observations are consistent with the efficient coding account if we assume that the visual system can adapt to environmental regularities over the course of evolution, resulting in feature biases that are encoded in the genome.

However, factors that are independent of visual input statistics may also separately contribute to the presence of cardinal orientation biases in animals. For example, some anatomical properties of the visual system naturally give a privileged status to the cardinal axes, such as the horizontal raphe of the retina, the role of the horizontal axis in vestibular and oculomotor system organization, and the distinction between processing of vertical and horizontal disparity

(Westheimer, 2003). Such properties need not be related to the orientation content of natural images, but may instead reflect general physical and/or developmental constraints. It is plausible that the presence of these architectural factors leads to cardinal biases, independent from the statistics of natural images. Thus, while past computational work suggests a strong correlational link between the statistics of the environment and the representation of orientation, the causal link between these observations has not yet been established.

Here, we use a causal manipulation to evaluate whether the efficient coding mechanism alone can account for the emergence of the oblique effect.

Specifically, we achieved this by measuring orientation representations in a convolutional neural network (CNN). We focus on the popular VGG-16 model, a standard feedforward network that achieves high performance at classifying objects in natural images (Simonyan & Zisserman, 2014). We first test whether a pre-trained VGG-16 model exhibits the classical oblique effect, assessed using the Fisher information measured at entire layers of the network, and the distribution of single-unit tuning properties. Assuming a definition of efficient coding in which mutual information between the stimulus and the network’s responses is maximized, we predict that Fisher information will be proportional to the square of the prior distribution (Figure 1D; Ganguli & Simoncelli, 2011; Wei & Stocker, 2015). In addition to a test of the efficient coding hypothesis, measuring orientation bias in this pre-trained model will provide an assessment of whether existing CNNs, often used as models of the primate visual system (Cichy & Kaiser, 2019; Kell & McDermott, 2019), exhibit this defining characteristic of biological vision.

**Figure 1.**
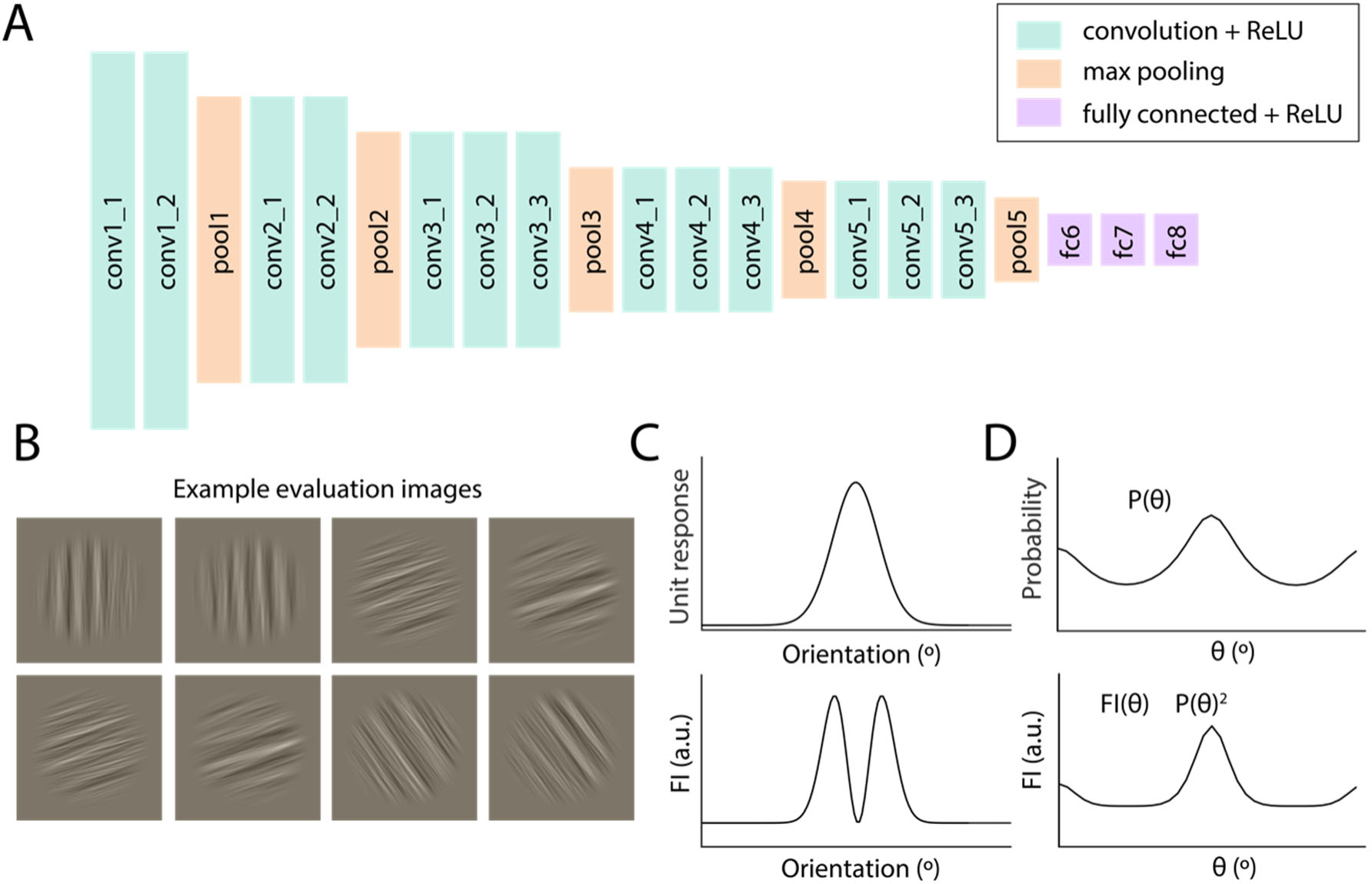
Evaluating orientation discriminability in a trained neural network model. **(A)** Schematic of the VGG-16 network architecture, with layers arranged from shallowest (left) to deepest. **(B)** Examples of oriented images used to measure orientation representations in the pre-trained network. Images were generated by filtering ImageNet images within a narrow orientation range, preserving their broadband spatial frequency content. Orientations varied between 0-179°, in steps of 1° (see *Methods, Evaluation stimuli)*. **(C)** Cartoon depiction of the approximate relationship between an example single unit tuning function and the Fisher information (FI) measured from that unit as a function of orientation. **(D)** Hypothetical depiction of the relationship between the prior distribution P over orientation θ, and the Fisher information (FI) when mutual information is maximized (Ganguli & Simoncelli, 2011; Wei & Stocker, 2015)

We next trained VGG-16 models on modified versions of the ImageNet database (Deng et al., 2009) that had been rotated by 0°, 22.5° or 45° relative to upright. This allowed us to determine whether a bias centered around other axes can be induced equally as well as a cardinal bias, and whether the biases observed in the pre-trained network were simply artifacts of some intrinsic property of the CNN (e.g. a square pixel grid that results in a cardinal reference frame). We demonstrate that, contrary to this alternative, networks trained on rotated images exhibited rotated biases that were consistent with the networks’ training set statistics. These results suggest that general visual experience with a non-uniform orientation distribution is sufficient to promote the formation of biased orientation representations. Further, our findings highlight how biased training data can fundamentally impact visual information processing in neural network models.

## Materials and Methods

### Training stimuli

During training, each model was presented with a modified version of the ILSVRC-2012-CLS training image set, a set of ∼1.3 million colored images with substantial variability in layout and background, each including an object in one of 1,000 categories (Deng et al., 2009; Russakovsky et al., 2015). Three modified versions of this image set were generated, corresponding to rotations of 0°, 22.5°, and 45° counter-clockwise relative to vertical. The purpose of generating a 0° (no rotation) version of the image set was to provide a control to isolate the effect of image rotation from any other properties of our modified image set.

To generate each version of the image set, we loaded each image from the original ILSVRC image set, rotated it by the specified amount, and cropped the image centrally by a specified amount that was the same for all rotations. Images were then scaled to a size of [224 x 224] pixels, and multiplied by a smoothed circular mask. The smoothed mask set to background all pixels at a radius of more than 100 pixels from the center, retained all pixels at a radius of less than 50 pixels from the center, and applied a cosine function to fade out the intermediate pixels. Finally, the background pixels were adjusted to a grey color that closely matches the mean RGB value of the training ImageNet images (Simonyan & Zisserman, 2014). All image processing for training set images was done in Python 3.6 (Python Software Foundation, Wilmington DE) using the Python Imaging Library. For each training set, a corresponding validation set was generated using the same procedure, and this validation set was used to evaluate performance during training. When preprocessing the images for training and validation, we modified the procedure from Simonyan and Zisserman’s paper by skipping the random rescaling and random left-right flipping steps. The purpose of this was to preserve the original spatial frequency and orientation content of the images as closely as possible.

### Evaluation stimuli

Networks were evaluated using sets of images that had known orientation content but were variable in their spatial phase and frequency (Figure 1B). These images consisted of randomly sampled images from the ILSRVC-2012-CLS image set which were filtered to have a particular orientation content. Before filtering each image, we first rotated it by a randomly chosen value in the range of 0-179 degrees, then cropped it centrally to a square and scaled to a size of [224 x 224] as described above. This was done to prevent any dependencies between orientation and other low-level image properties, such as spatial frequency content and luminance contrast, in the final filtered images. After this step, we converted to grayscale, z-scored the resulting luminance values, and masked the image with the smoothed circular mask described above. The image was then padded with zeros to a size of [1012 x 1012] pixels, and transformed into the frequency domain (using *fft2.m*). We then multiplied the frequency-domain representation by an orientation filter and a spatial frequency filter. The orientation filter consisted of a circular Gaussian (Von Mises) function centered at the desired orientation, with concentration parameter (k) of 35 (full-width at half-max=11.5°). The spatial frequency filter was a bandpass filter from 0.02 to 0.25 cycles/pixels, with Gaussian smoothed edges (smoothing SD=0.005 cycles/pixel). After multiplying by these filters, we then replaced the image’s phase with random values uniformly sampled between -pi to +pi (to randomize the spatial phase of oriented elements in the image) and transformed back into the spatial domain (using *ifft2.m*). Next, we cropped the image back to its original size of [224 x 224], multiplied again by the smoothed circular mask, and converted the image into a 3-channel RGB format. Finally, the luminance in each color channel was normalized to have a mean equal to the mean of that color channel in the training ImageNet images and a standard deviation of 12 units. All image processing for the evaluation image sets was done using Matlab R2018b (MathWorks, Natick MA).

Using the above procedures, we generated four evaluation image sets, each starting with a different random set of ImageNet images. Images in each evaluation set had orientations that varied between 0° and 179°, in steps of 1°, resulting in 180 discrete orientation values. Throughout this paper, we use the convention of 0° for vertical and 90° for horizontal orientations, with positive rotations referring to the clockwise direction, and negative rotations referring to the counter-clockwise direction. Each evaluation set included 48 examples of each orientation, for a total of 8640 images per set.

### Measuring image set statistics

To verify that the modified versions of the ImageNet images had the anisotropic orientation statistics that we expected, we measured the orientation content of each training image using a Gabor filter bank. The filter bank included filters at orientations from 0° to 175° in 5° steps, at spatial frequencies of 0.0200, 0.0431, 0.0928, and 0.200 cycles per pixel (orientation bandwidth of filters was 19°). The filter bank was generated using the *gabor.m* function in Matlab R2018b (MathWorks, Natick MA). Since all filtering was performed in the Fourier domain, we also used a custom modified version of the *gabor.m* function which allowed us to directly generate a frequency-domain representation of each filter (Jain & Farrokhnia, 1991). Before filtering each image, we converted it to grayscale, subtracted its background color so that the background was equal to zero, and padded each image with zeros to a size of 1012 x 1012 pixels (this was the size needed to accommodate the lowest frequency filter). Images were then converted into the frequency domain for filtering (using *fft2.m*), and multiplied by the filter bank. Next, we converted back to the spatial domain and un-padded the image back to its original size (224 x 224 pixels). Finally, we took the magnitude of the filtered image, and averaged the magnitude across all pixel positions to obtain a single value for each filter orientation and spatial frequency. Next, for each image, within each spatial frequency, we converted the orientation magnitude values into an estimated probability distribution by dividing by the sum of the magnitude across all orientations. Since this was done for all orientations of one spatial frequency at a time, this correct for differences in power across spatial frequency and facilitates combining results across spatial frequency. Results were similar within each spatial frequency individually; we averaged over spatial frequency to produce the final plots (Figure 6B). This analysis was done on the training set images only, which included ∼1300 images in each of 1000 categories, for a total of ∼1.3 million images.

### Network training and evaluation

We trained VGG-16 networks (Simonyan & Zisserman, 2014) on three different modified versions of the ImageNet dataset (see *Training stimuli* for details). For each of the three image sets, we initialized and trained four VGG-16 networks (replicates), giving a total of 12 models. All models were trained using Tensorflow 1.12.0 (Abadi et al., 2016), using the TF-slim model library (Silberman & Guadarrama, 2016) and Python 3.6 (Python Software Foundation, Wilmington DE). All models were trained using the RMSProp algorithm with momentum of 0.80 and decay of 0.90. The learning rate was 0.005 with an exponential decay factor of 0.94, and the weight decay parameter was 0.0005. Networks were trained until performance on the validation set (categorization accuracy and top-5 recall) began to plateau, which generally occurred after around 350K-400K steps. The validation images used to evaluate performance were always rotated in an identical manner to the training set images. Training was performed on an NVIDIA Quadro P6000 GPU (NVIDIA, Santa Clara CA). All evaluation was performed using the first checkpoint saved after reaching 400K steps. As noted above, we did not perform data augmentation steps during image pre-processing for training. Removing these procedures may have contributed to the relatively low final classification performance that we observed (top-5 recall accuracy ∼60%).

To measure activations from each trained network, we split the evaluation image sets (consisting of 8640 images each) into 96 batches of 90 each. We then passed each batch through each trained network and measured the resulting activations of each unit as the output of the activation function (a rectified linear operation). We saved the activations for each unit in each layer for all images, which were then submitted to further analysis. We performed this evaluation procedure on a total of 17 networks: the 12 models trained on modified ImageNet images, a pre-trained VGG-16 network from the TF-slim model library (Silberman & Guadarrama, 2016), and four randomly initialized VGG-16 models that served as a control. All subsequent analyses were performed using Python 3.6 (Python Software Foundation, Wilmington DE).

### Computing Fisher information (FI)

To measure the ability of each network layer to discriminate small changes in orientation, we estimated Fisher information (FI) as a function of orientation. To estimate FI for each network layer, we first computed FI for each unit in that layer, then combined information across units. FI for each unit was computed based on the slope and variance of that unit’s activation at each point in orientation space, according to the following relation:

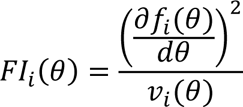

Where *f*_*i*_(*θ*) is the unit’s measured orientation tuning curve, and *v*_*i*_(*θ*) is the variance of the unit’s responses to the specified orientation. We estimated the slope of the unit’s tuning curve at *θ* based on the difference in its mean response (*μ*_*i*_) to sets of images that were Δ=4° apart (using different values of Δ did not substantially change the results).

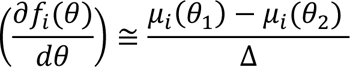

Where

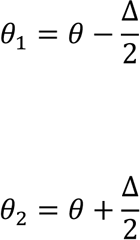

We presented an equal number of images (48) at each orientation, so the pooled variance was calculated as:

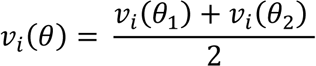

Finally, we summed this measure across units of each layer to obtain a population level estimate of FI. Note that this measure does not account for the covariance among units, see *Multivariate analyses* for complementary approaches.

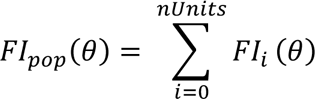

Where *nUnits* is the number of units in the layer. We computed *FI*_*pop*_(*θ*) for theta values between 0° and 179°, in steps of 1°. When plotting FI, to aid comparison of this measure across layers with different numbers of units, we divided *FI*_*pop*_ by the total number of units in the layer, to capture the average FI per unit. We note that this analysis was performed across all units at each layer, not excluding any units whose spatial receptive field was outside the circular stimulus region. These non-responsive units contributed zero to the Fisher information sum at early layers. We note also that due to the different numbers of units per layer, the absolute values of FI are not directly comparable across layers.

### Multivariate analyses

In addition to calculating the sum of univariate Fisher information across all individual units at each network layer (see previous section, *Computing Fisher information)* we were also interested in evaluating the information content of multivariate patterns of activation across entire layers of the network. To this end, we computed a multivariate version of Fisher information, using the following expression (Abbott & Dayan, 1999):

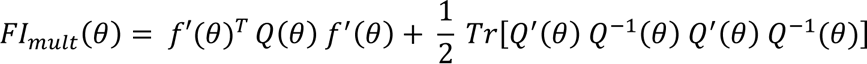

Where *Q*(*θ*) is the pooled covariance matrix, computed as:

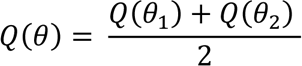

And *Q*′(*θ*) was the estimated derivative of the covariance matrix, obtained as:

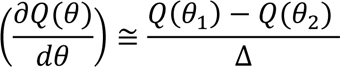

Where *θ*_1_ and *θ*_2_ are as defined in the previous section. *Tr* denotes the trace operation, and *Q*^-1^(θ) is the inverse of the covariance matrix. Values of *θ* and *f*′(*θ*) were as defined in the previous section.

To make computing the covariance matrix computationally feasible, we first performed principal components analysis on the activation matrix from each network layer (for each image set, a matrix of size [8640 *x nUnits*]), implemented using Scikit-learn in Python 3.6 (Python Software Foundation, Wilmington DE). We then computed the above expression for Fisher information using the scores for the top N principal components (PCs), for values of N ranging from 2 to 47 (there were 48 images per orientation, so covariance matrix estimates became unstable when using more features). Values were similar for different values of N (several examples of varying N shown in Figure 4B).

In addition to computing multivariate Fisher information in this reduced-dimensionality space, we used the principal components to estimate the dimensionality of the orientation representations at each layer (Figure 5C). The purpose of this was to facilitate qualitative comparisons of the representational structure across layers. To isolate the dimensions that were related to orientation, and not to variability across images having the same orientation, we first averaged across images of the same orientation (e.g. collapsing over any variability not related to orientation). This resulted in a matrix of size [180 x N], where N is the total number of units in the layer. Next, we performed PCA on this averaged matrix, and calculated the cumulative variance explained by each principal component. Our estimate of dimensionality was the PC number after which additional components contributed less than 5% additional variance. This threshold is somewhat arbitrary but captures the approximate point at which an “elbow” appears in a plot of cumulative variance versus PC number (Figure 5C). We note that this estimate of dimensionality is not exact and is not cross-validated, but should provide a reasonable index of how dimensionality changes across model layers.

Finally, we computed an additional measure of multivariate orientation separability in the reduced-dimensionality PC space. The rationale for this was to provide a complementary measure to the Fisher information and ensure that the results we obtained with FI were not specific to that method. For this measure, as with FI, we computed a statistic for each point *θ* in orientation space, using the responses to two orientations Δ=4° apart (as before, changing Δ did not substantially change the results). This resulted in two “clouds” of points in N-dimensional PC space, corresponding to the two orientations (see Figure S2). We first exhaustively computed the Euclidean distances between all pairs of points in different clouds (48^2 = 2304 total distances). Next, we computed a t-statistic for these distances: the mean of all distance values divided by the standard deviation of all distance values. This measure reflects the reliability of the separation between point clouds corresponding to different orientations. We computed this measure for several different values of N.

### Fisher information bias (FIB)

To quantify the amount of bias (non-uniformity) in Fisher information at each layer of the network, we computed a measure which we refer to as the Fisher information bias (FIB). For the pre-trained model and the networks trained on upright images, we expected the network to over-represent cardinal orientations, showing peaks in FI around vertical and horizontal.

However, the models trained on rotated images were expected to show peaks rotated by a specified amount relative to the cardinal orientations. To account for these different types of bias, we computed three versions of the FIB: one that measures the height of peaks in FI around the cardinal orientations (FIB-0), one that measures the height of peaks in FI that are 22.5° counter-clockwise of the cardinals (FIB-22), and one that measures the height of peaks in FI that are 45° counter-clockwise of the cardinals (FIB-45), relative to a baseline. The equation for each FIB measure is as follows:

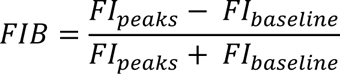

Where *FI*_*peaks*_ is the sum of the FI values in a range ±10° around the orientations of interest (0° and 90° for FIB-0, 67.5° and 157.5° for FIB-22, and 45° and 135° for FIB-45), and *FI*_*baseline*_ is the sum of the FI values in a range ±10° around the orientation chosen as a baseline (22.5° and 112.5°). Since FI is necessarily positive, each of these FIB measures can take a value between +1 and -1, with positive values indicating more information near the orientations of interest relative to the baseline (peaks in FI), and negative values indicating less information near the orientations of interest relative to baseline (dips in FI). An analogous method was used to compute the bias in multivariate FI (Figure 4) as well as the multivariate t-statistic (Figure S2; see previous section, *Multivariate analyses*).

To test whether FIB differed significantly between trained models and the randomly initialized (not trained) models, we performed non-parametric t-tests between FIB values corresponding to each training set and the random models. Specifically, we tested the hypothesis that the primary form of bias measured in models corresponding to each training set (e.g. FIB-0 for the models trained on upright images, FIB-22 for the models trained on 22.5° rotated images, FIB-45 for the models trained on 45° rotated images) was significantly higher for the models trained on that image set than for the random (not trained) models. Since we generated four replicate models for each training image set, and evaluated each model on four evaluation image sets, there were 16 total FIB values at each layer corresponding to each training set. To compare the FIB values corresponding to each training set against the random models, we first calculated the “real” difference in FIB between the groups, based on comparing the mean of the 16 values for the trained models versus the mean of the 16 values for the random models. Next, we concatenated the values for the trained and random models (32 values total) and randomly shuffled the group labels across all values 10,000 times. For each of these 10,000 shuffles, we then computed the difference between groups based on the shuffled labels (“shuffled” differences). The final p-value was generated by calculating the number of iterations on which the shuffled difference exceeded the real difference and dividing by the number of total iterations. The p-values were FDR corrected across model layers at q=0.01 using SciPy (Benjamini & Yekutieli, 2001). The same procedure was used to test for differences in FIB-0 between the pre-trained model and the control model (note that there was only one replicate for the pre-trained model, so this test included only 4 data points per condition).

### Single-unit tuning analysis

To measure the extent to which training set statistics impacted the orientation tuning of individual units in each network, we measured tuning functions based on each unit’s responses to the evaluation image set, and we performed curve fitting to quantify tuning properties. First, we measured an orientation tuning function for each unit at each layer of the model by averaging its responses to all evaluation set images that had the same orientation (in each image set, there were 48 images at each of 180 orientations). Any units that had a constant response across all images or a zero response to all images were removed at this stage (this included mainly units whose spatial selectivity was outside the range stimulated by the circular image aperture, around 35% of units per layer at the earliest layers). We computed and saved an orientation tuning curve for each unit in response to each of the four evaluation image sets. We then averaged over these four evaluation sets before fitting.

To characterize the tuning curves, we fit each with a circular Gaussian (Von Mises) function, having the basic form:

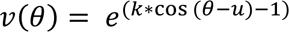

Where *u* is a parameter that describes the center of the unit’s tuning function, and *k* is a concentration parameter that is inversely related to the width of the tuning function. In this formulation, the *k* parameter modifies both the height and the width of the tuning function. To make it possible to modify the curve’s height and width independently, we normalized the Von Mises function to have a height of 1 and a baseline of 0, and then added parameters for the amplitude and baseline, as follows:

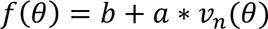

Where *v*_*n*_(*θ*) denotes the Von Mises function after normalization. This resulted in a curve with four total parameters: center (*u*), concentration parameter (*k*), amplitude, and baseline.

We fit a curve of this form to each unit’s average tuning function using linear least-squares regression, implemented with the optimization library in SciPy (version 1.1.0). To initialize the fitting procedure, we used the argmax of the tuning function as an estimate of its mean, the minimum value as an estimate of its baseline, and the range as an estimate of its amplitude. The concentration parameter k was always initialized at 1. Values for the center were constrained to lie within the range of [-0.0001, 180.0001], k was constrained to positive values >10^-15^, and amplitude and baseline were allowed to vary freely. To prevent any bias in the center estimates due to the edges of the allowed parameter range, we circularly shifted each curve by a random amount before fitting.

After fitting was complete, we assessed the goodness of the fit using R^2^. To assess the consistency of tuning across different versions of the evaluation image set, we used R^2^ to assess the fit between the single best-fit Von Mises function (computed using the tuning function averaged over all evaluation image sets) and each individual tuning curve (there were four individual tuning curves, each from one version of the evaluation image set). We then averaged these four R^2^ values to get a single value. We used a threshold of average R^2^ > 0.40 to determine which units were sufficiently well-fit by the Von Mises function, and retained the parameters of those fits for further analysis.

### Code Availability

All code is available on the author’s Github page (link to be added upon publication).

## Results

We measured the activation responses of several trained VGG-16 networks (Figure 1A) (Simonyan & Zisserman, 2014) presented with oriented images (Figure 1B) to evaluate whether each network showed non-uniformities in its orientation representations across feature space. First, we tested whether a pre-trained VGG-16 model (Silberman & Guadarrama, 2016) exhibits the classical oblique effect. Next, we evaluated whether this bias changed in a predictable way when networks with the same architecture were trained on modified versions of the ImageNet database (Figure 6A).

### Measuring cardinal biases in a pre-trained VGG-16 model

We first evaluated non-uniformity at the level of each pre-trained network layer by computing the layer-wise Fisher information (FI), which reflects how well each layer’s activations can distinguish small changes in orientation (see *Methods, Computing Fisher information*). Briefly, the contribution of each network unit to the layer-wise FI is the squared slope of a unit’s tuning function at each orientation normalized by the variance of the response at that orientation. Thus, the steep part of a unit’s tuning function will carry more information because two similar orientations will evoke different responses (Figure 1C). However, the flat parts of a unit’s tuning curve (i.e. at the peak or in the tails) will not carry very much information because the unit will respond about the same to two similar orientations. We focused on the Fisher information because it has been suggested to have a predictable relationship to the prior orientation distribution. Specifically, when mutual information is maximized, Fisher information is predicted to be proportional to the square of the prior distribution (Figure 1D; Ganguli & Simoncelli, 2011; Wei & Stocker, 2015). This therefore provides a testable prediction of the efficient coding account. To introduce the variability necessary for calculating Fisher information, we used orientation bandpass-filtered natural images which create stimulus-level variability in the images for a given orientation.

For a pre-trained VGG-16 model, plotting FI as a function of orientation reveals noticeable deviations from uniformity, particularly at deep layers of the network (navy blue curves in Figure 2A). While the first layer of the model (conv1_1), gives a relatively flat profile of FI with respect to orientation, by layer 7 (conv3_1), peaks in FI are apparent around the cardinal orientations, 0°/180° and 90°. At later layers of the model, the peaks in FI are more pronounced and begin to take on a characteristic double-peaked shape, where FI is maximal slightly (∼5°) to the left and right of the cardinal orientations, with a dip at the exact position of the cardinal orientations (this shape is discussed in more detail in the next section after we report statistics about the preferred orientation and width of single unit tuning functions). In contrast, when the same analysis is done on a randomly initialized VGG-16 model (no training performed), FI is flat with respect to orientation at all layers, suggesting that a randomly initialized model does not exhibit this same cardinal bias (gray curves in Figure 2A).

**Figure 2.**
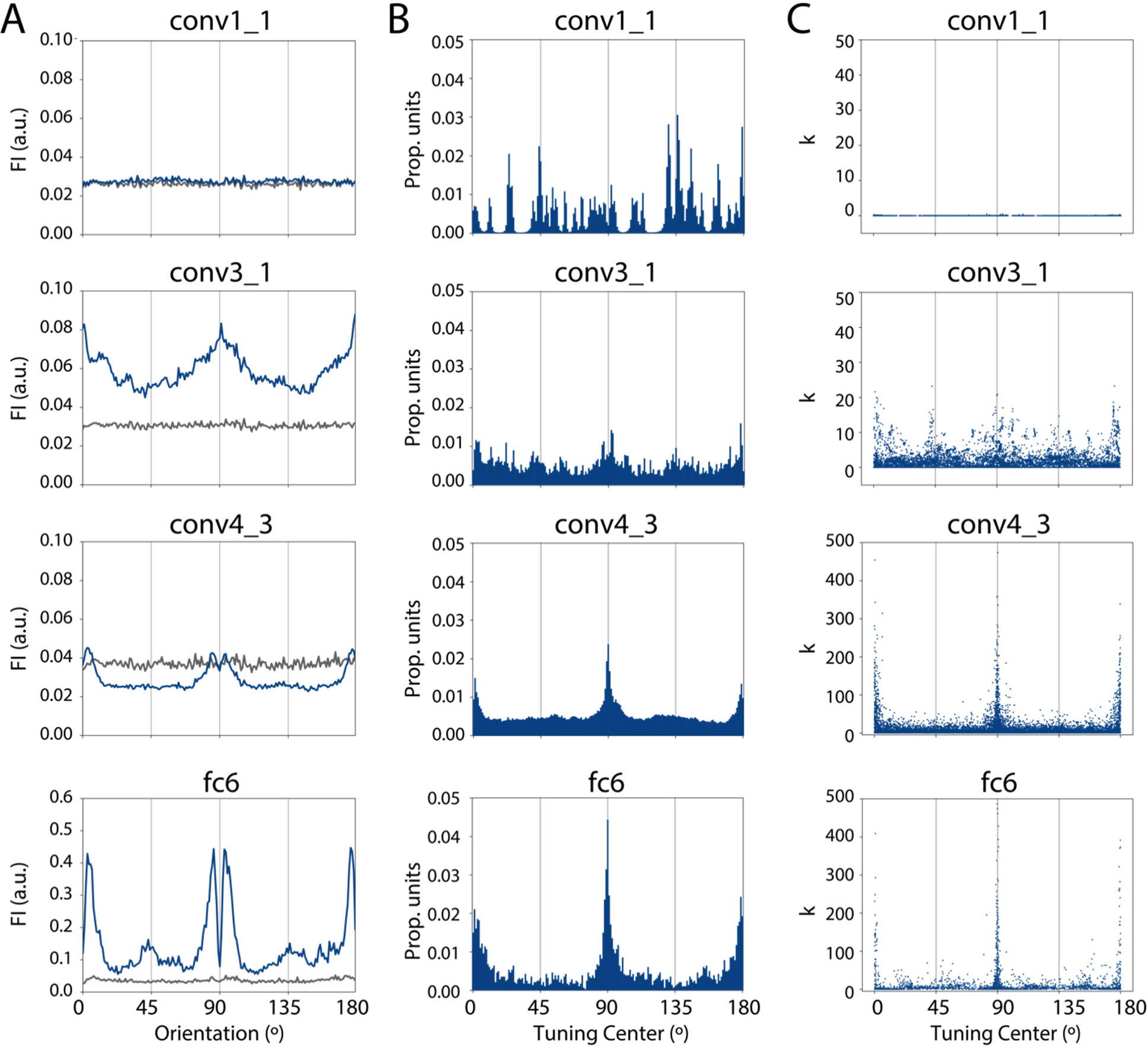
Pre-trained VGG-16 shows maximum orientation information just off of cardinal orientations, and non-uniformity in the distribution of single unit tuning properties. **(A)** FI is plotted as a function of orientation for several example layers of the pre-trained model (navy blue) and a randomly initialized model (gray). See *Methods, Computing Fisher information* for details. **(B)** Distribution of the tuning centers of pre-trained network units that were well-fit by a Von Mises function. See Figure S1 for the proportion of well-fit units per layer, and the distribution of centers for the randomly initialized model. **(C)** Concentration parameter (k) versus center for individual units in the pre-trained model (data in the top three panels of C have been down-sampled to a maximum of 10,000 points for visualization purposes).

To quantify this effect at each layer, we computed a metric which we term the Fisher Information Bias (FIB), which captures the relative height of the peaks in FI compared to a baseline (see *Methods, Fisher information bias*). We defined three versions of this metric, the FIB-0, FIB-22, and FIB-45, which denote the height of peaks in FI around the cardinal orientations, around orientations 22.5° counter-clockwise of cardinals, and around orientations 45° counter-clockwise of cardinals, respectively. For example, to get the FIB-0, we take the mean FI in 20° bins around 0° and 90°, subtract the mean FI in in a baseline orientation range, and divide by the sum of these two means. Because the pre-trained model showed peaks in FI around cardinals only, we focus on the FIB-0 in this section; the FIB-22 and FIB-45 are discussed in the following section (*Training networks on rotated images*). We found that for the pre-trained model, the FIB-0 increased with depth in the network, showing values close to zero for the first four layers, then showing positive values that increase continuously at each layer (navy blue line in Figure 3). In contrast, we found less evidence for a cardinal bias in the randomly initialized model, shown by smaller values of the FIB-0 at all layers (gray line in Figure 3). The difference in FIB-0 between the pre-trained and randomly initialized models was significant starting at the fifth layer (conv2_2), and at all layers deeper than conv2_2 (one-tailed non-parametric t-test, FDR corrected q=0.01). However, there was a small increase in the FIB-0 at the later layers of the randomly initialized model, reflecting a weak cardinal bias (at the deepest layer, the FIB-0 was still more than 5x as large for the pre-trained model as for the random model). We return to this issue for more consideration in the *Discussion*.

**Figure 3.**
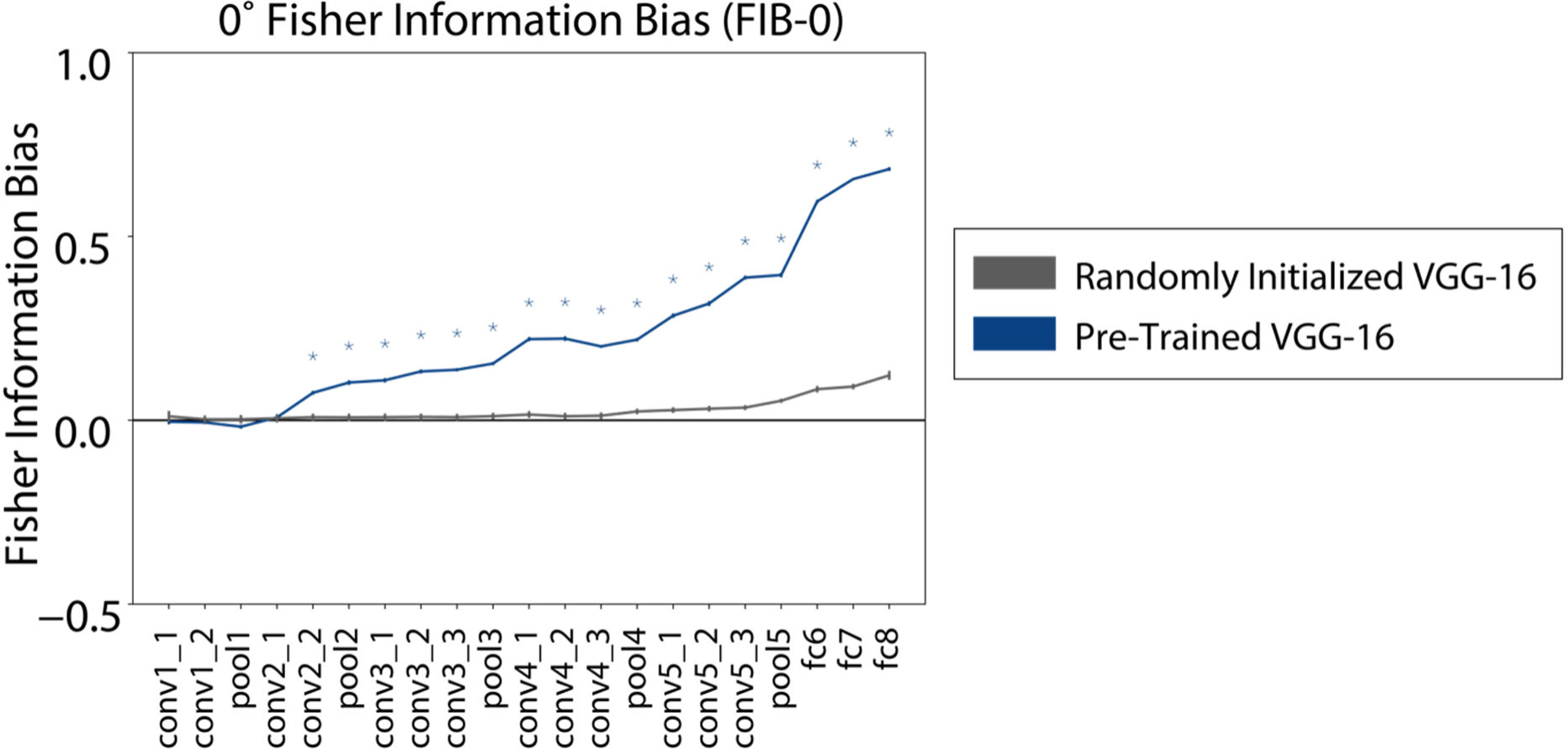
Cardinal bias in a pre-trained VGG-16 model increases with depth. FIB-0, a measure of cardinal information bias (see *Methods, Fisher information bias)*, plotted for a pre-trained model (navy blue) and a randomly initialized control model (gray), with asterisks indicating layers for which the pre-trained model had significantly higher FIB-0 than the random model (one-tailed nonparametric t-test, FDR corrected q=0.01). Error bars reflect standard deviation across four evaluation image sets.

Having demonstrated that a pre-trained CNN exhibits an advantage for discriminating cardinal versus other orientations, we were next interested in whether this bias was linked to the distribution of tuning properties across single units at each layer, as has been observed in the brains of animals such as cats and macaques (Li et al., 2003; Shen et al., 2014; Vogels & Orban, 1994). To investigate this, we computed the average orientation tuning profiles for individual units in response to stimuli of all orientations and fit these profiles with Von Mises functions to estimate their center and concentration parameter (or width, denoted *k*). Units that were not well-fit by a Von Mises were not considered further (approximately 30% of all units, see *Methods, Single-unit tuning analysis* and Figure S1. Figure 2B shows the distribution of fit centers for all units in four example layers of the pre-trained model that were well-fit by a Von Mises function. These distributions show peaks at random locations for the first layer of the network, but exhibit narrow peaks around the cardinal orientations for the deeper conv4_3 and fc6 layers. In contrast, the randomly initialized model did not show an over-representation of cardinal-tuned units (Figure S1). In addition, plotting the concentration parameter for each unit versus the center (Figure 2C) shows that for the deepest three layers shown, the most narrowly-tuned units (high k) generally have centers close to the cardinal orientations.

Together, these findings indicate that middle and deep layers of the pre-trained network have a large proportion of units tuned to cardinal orientations, and that many of these units are narrowly tuned.

These findings may provide an explanation for the double-peaked shape of the FI curves for the pre-trained model at deep layers (Figure 2A). Since FI is related to the slope of a unit’s tuning function, it is expected to take its maximum value on the flanks of a tuning curve, where slope is highest, and take a value of zero at the tuning curve peak (Figure 1C). Thus, having a large number of narrowly-tuned units with their peaks precisely at 0° and 90° could result in layer-wise FI having local maxima at the orientations just off of the cardinals. We note also that these findings indicate a divergence from the predicted shape of FI based on mutual information maximization (Figure 1D); we return to this issue for further consideration in the *Discussion*.

In addition to the above analyses, we were also interested in determining whether biases were present in activation patterns across all units in each layer. To achieve this, we computed the multivariate Fisher information (see *Methods, Multivariate analyses*). Note that this is distinct from the version of FI described earlier in this section (Figure 2, Figure 3), which combines information linearly across units but does not take advantage of the covariance structure of the data. Before calculating multivariate FI, we first reduced the dimensionality of the data using principal components analysis (PCA; see *Methods, Multivariate analyses).* As shown in Figure 4, multivariate FI revealed a similar pattern of results as the summed univariate FI discussed previously. FI was generally flat across orientation space for the earliest layers, and peaks around the cardinal orientations began to emerge at the middle layers (around conv3_1, for this analysis). As with the univariate version of FI, multivariate FI at the deeper layers of the model exhibited a pronounced double-peaked shape with highest values a few degrees clockwise and counter-clockwise of the cardinal orientations. Furthermore, when we utilized another measure of orientation discriminability in PC space, based on the reliability of distances between points corresponding to nearby orientations (Figure S2, see *Methods, Multivariate analyses)*, we again found biases that were similar in form and emerged at middle layers of the model. In contrast to FIB, the cardinal bias in this measure did not exhibit a continual increase with depth in the model, however it was above zero for all middle and late layers. Together, these results suggest that the presence of pattern-level orientation biases in VGG-16, and the fact that these biases persist until the deepest model layers, is not dependent on the specific information metric used.

**Figure 4.**
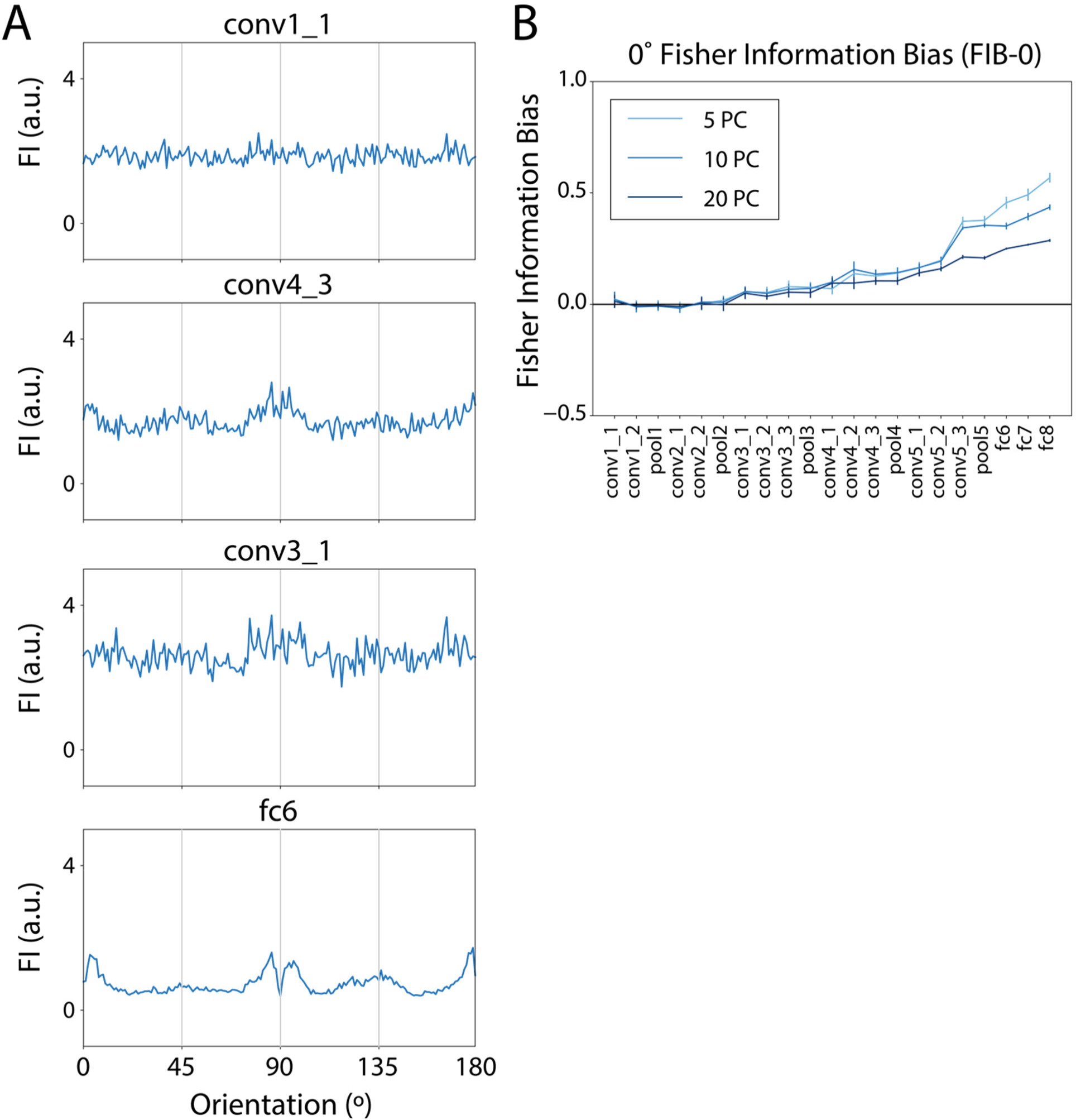
Multivariate FI shows similar pattern of results as summed univariate FI. **(A)** Multivariate Fisher information, calculated after performing PCA, shown for four example layers in the pretrained model. See methods (*Multivariate analyses*) for details. This figure reflects the calculation with first 10 PCs retained; similar results were found with different numbers of PCs retained. **(B)** Multivariate version of FIB-0 is plotted for each layer, comparing results with 5, 10 or 20 principal components retained. Error bars reflect ±1 standard deviation of the measure across 4 evaluation image sets.

The above univariate and multivariate analyses suggest that the structure of orientation representations differs at early, middle, and deep layers of VGG-16. To further explore this difference and visualize the representations at each layer, we plotted the representations of images in the space spanned by the first two principal components (Figure 5A). We also plotted the orientation tuning profiles of the first four principal components of several example layers (Figure 5B). This revealed that at the earliest layer of the model (conv1_1), the representation of orientation appeared to be well-described by two principal components resembling a sine and cosine function, resulting in an approximately circular representation in PC space. The third and fourth principal components for this layer showed little response change as a function of orientation. In contrast, the middle and deeper layers of the model exhibited more complex response profiles. At conv3_1 and conv4_3, PCs 1 and 2 were similar to those of the first layer, but the third and fourth principal components tended to be higher frequency or include sharp points near the cardinal orientations. At a deep fully-connected layer (fc6), the tuning profiles were no longer sinusoidal, and instead exhibited steep points and/or dips near the cardinal axes. The steepest regions of these principal component tuning profiles appear to coincide with the peaks we measured in FI (Figure 2A, Figure 4), suggesting these response profiles may underly the strong biases we observed at deep layers.

**Figure 5.**
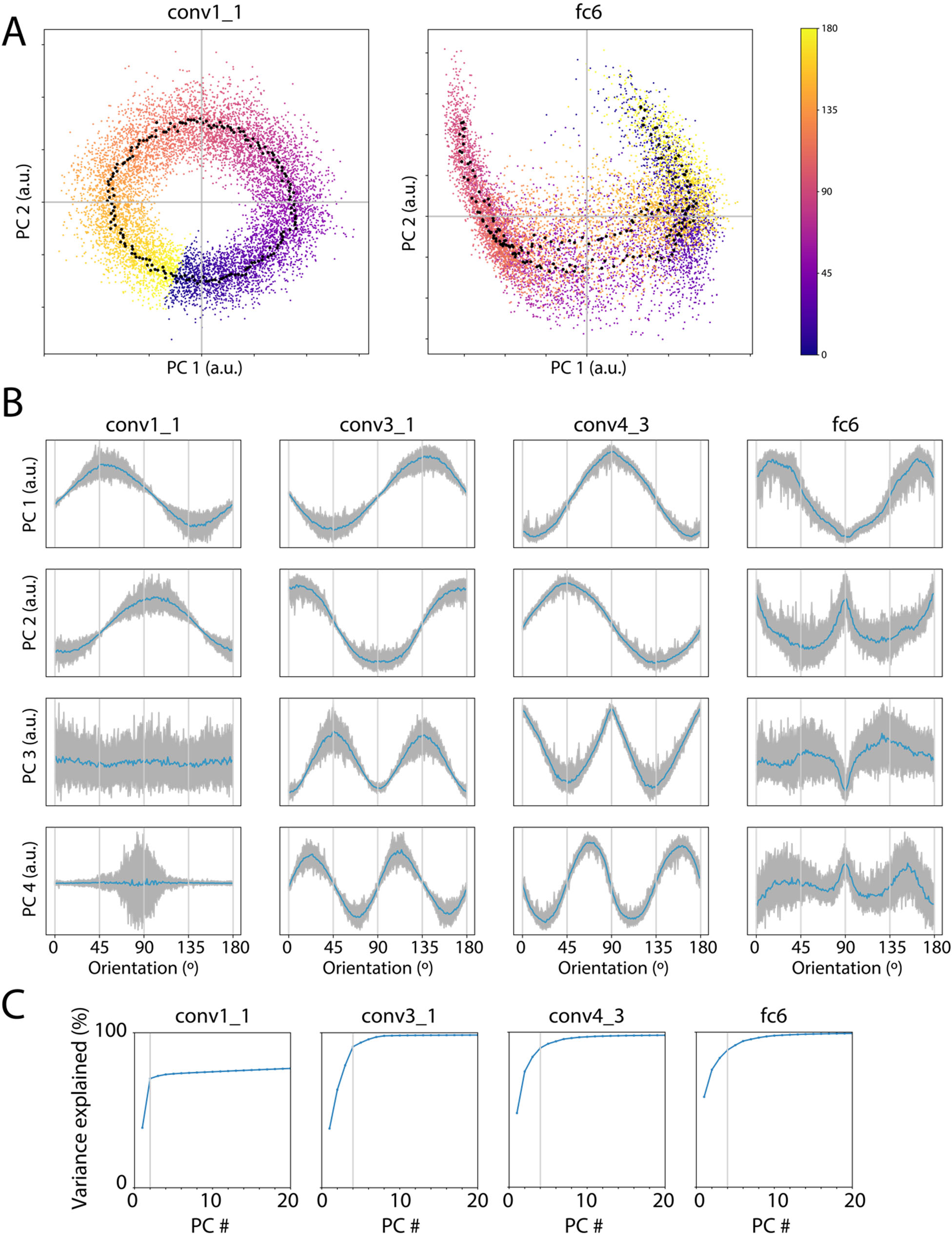
Principal component analysis reveals a graded change in the structure of orientation representations across pre-trained model layers. **(A)** For two example layers (conv1_1 and fc6), scatter plot of scores corresponding to the first two principal components of the layer’s representation (see *Methods, Multivariate analyses)*. Colored points indicate individual images, with color indicating stimulus orientation. Black points indicate the mean of the 48 points corresponding to each orientation. **(B).** Scores for the first four principal components are plotted as a function of orientation, for several example layers. Blue lines indicate the mean value of that principal component score as a function of orientation, gray lines indicate individual images. **(C)** Percent variance explained by each principal component of the data, after averaging across trials of a common orientation. Vertical line indicates the number of components after which additional components contribute <5% additional variance (see *Methods, Multivariate analyses*).

Additionally, these more complex response profiles may imply an increase in the dimensionality of the space spanned by all distinct orientations. To estimate the dimensionality of this subspace, we averaged over different images at each orientation, and performed PCA on the resulting matrix (see *Methods, Multivariate analyses)*. When we plot the cumulative percent variance explained by each component in this representation (Figure 5C), we find distinct patterns between early and late layers. While early layers exhibit a clear plateau where components after 2 contribute little additional variance, later layers exhibit a smoother function where more components are required to reach a plateau. Together, these observations support the idea that the dimensionality of orientation representations increases from shallow to deeper layers of VGG-16, with representations morphing from a veridical representation of the circular feature space spanned by orientation, to a more complex and biased format.

### Training networks on rotated images

Having demonstrated that a pre-trained VGG-16 network exhibits a much stronger cardinal orientation bias compared to a randomly initialized network, we next tested whether training a model on rotated images would result in rotated biases. This test is needed to demonstrate that the frequently-observed cardinal bias is not the only possible orientation bias that can be induced in a visual system through exposure to a set of images with non-uniform statistics. We trained networks on three modified versions of the ImageNet dataset (Deng et al., 2009), consisting of images that were rotated by either 0°, 22.5°, or 45° in a clockwise direction relative to the upright position (Figure 6A). Separately, we also verified that the image statistics of each of the modified sets exhibited the expected distribution, such that vertical and horizontal orientations were most common in the upright training set, orientations 22.5° counter-clockwise of cardinals were most common in the -22.5° rotated set, and orientations 45° counter-clockwise of cardinals were most common in the -45° rotated set (Figure 6B).

**Figure 6.**
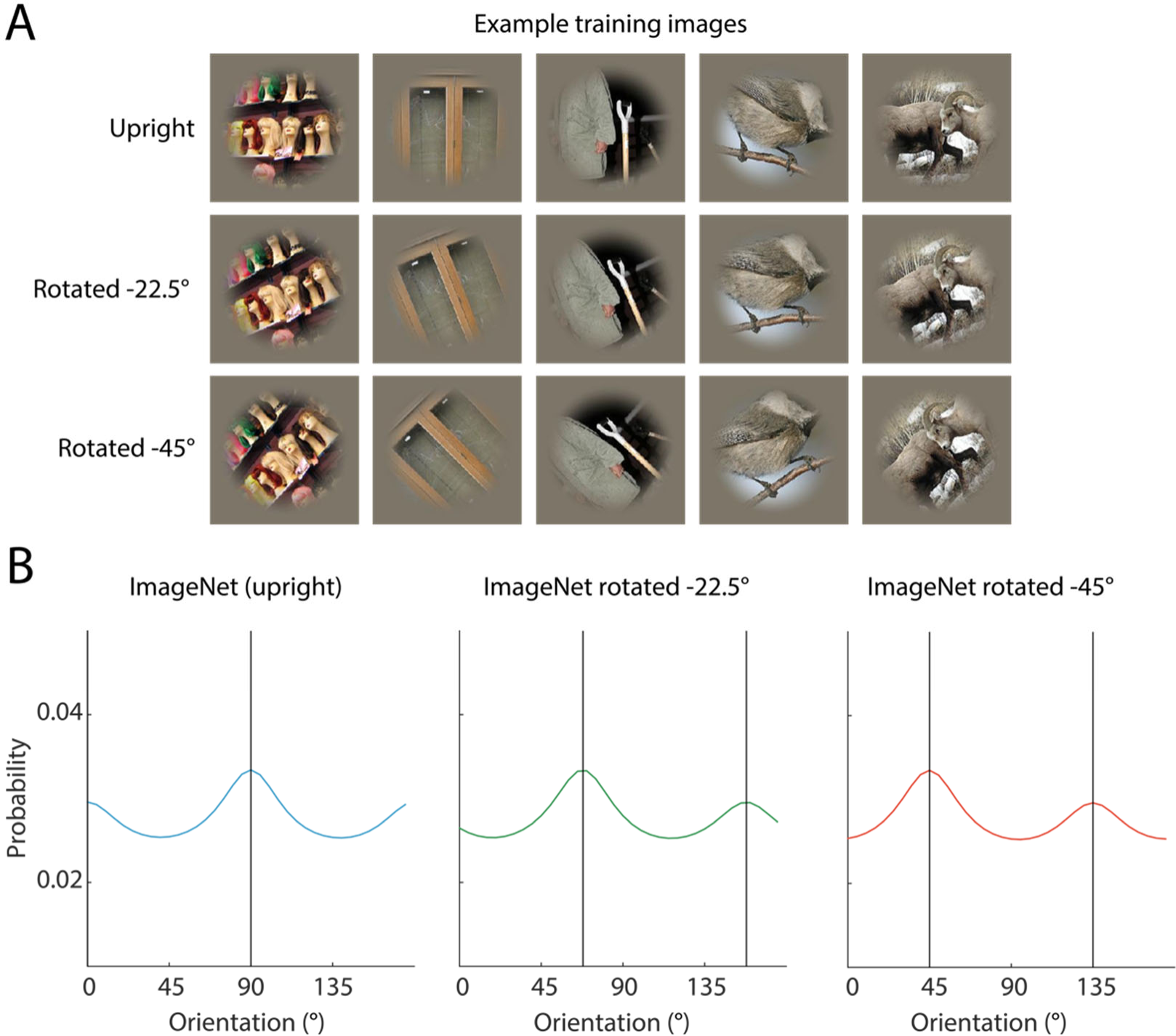
Rotated images used to train VGG-16 networks. **(A)** Separate networks were trained on either upright or rotated versions of the ImageNet image set, with a smoothed circular mask applied to remove vertical and horizontal image edges. **(B)** Orientation content from images in each of the training sets in (A) was measured using a Gabor filter bank (see *Methods, Measuring image set statistics*).

Given the hypothesized relationship between FI and the prior probability distribution (Figure 1D), we predicted that the FI curves for each of these models would have peaks closely aligned with the peaks of its respective prior distribution.

Our results indicate that training on rotated images shifted the orientation bias by a predictable amount. FI for the models that were trained on upright images shows a relatively similar shape to the pre-trained model, with peaks appearing at a few degrees to the left and right of the cardinal orientations (Figure 7A). This demonstrates that though our training procedure and image set were not identical to those used for the pre-trained model, they resulted in the formation of similar orientation biases. In contrast, the models trained on rotated images each showed a FI curve that was similar in shape but shifted relative to the curve from the model trained on upright images, such that the peaks in FI were always near the orientations that were most common in the training set images (Figure 7D,7G).

**Figure 7.**
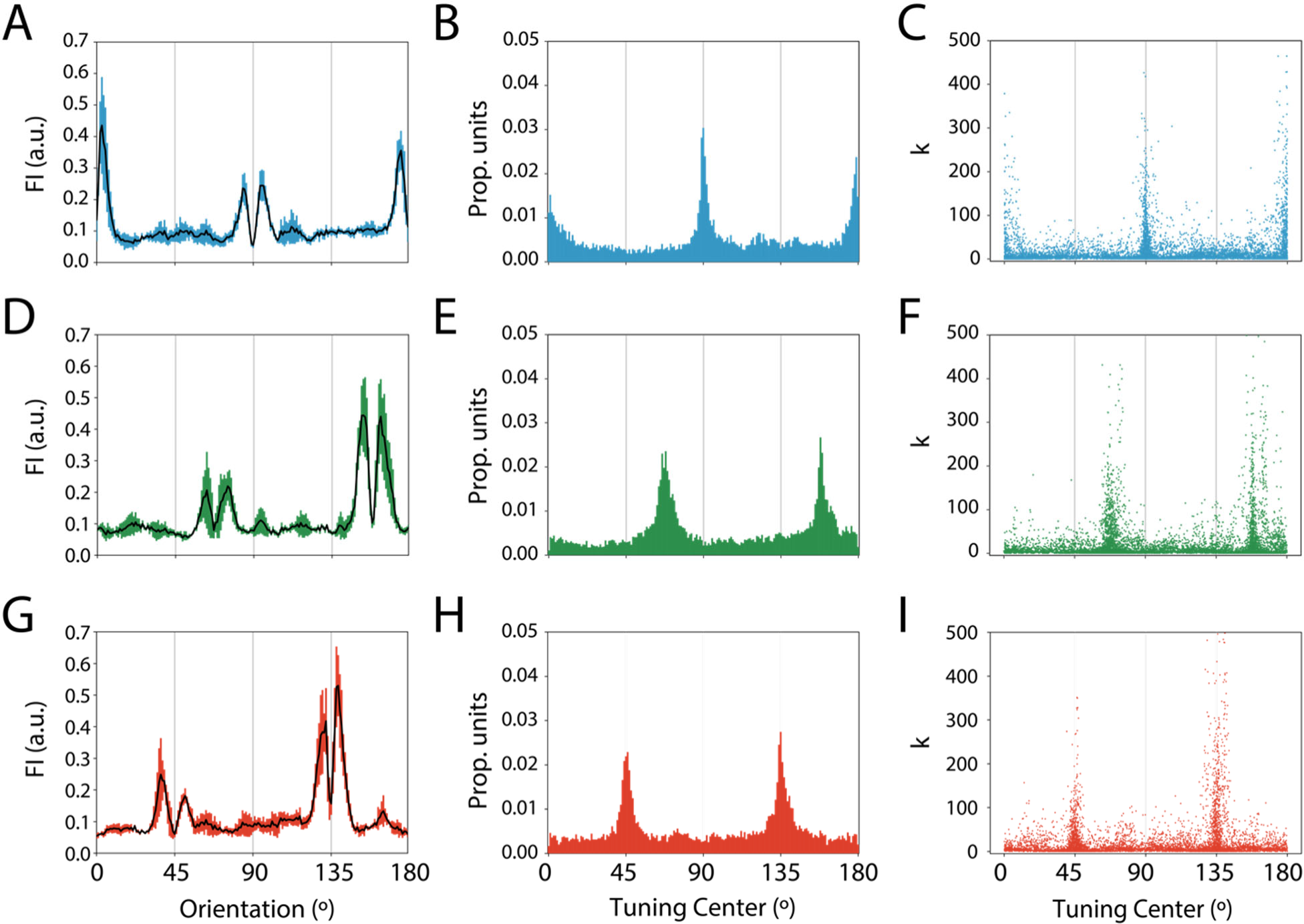
When networks are trained on rotated images, both population-level information and single unit tuning distributions reflect modified training set statistics. **(A-C)** show data from one example layer (fc6) of four separately initialized networks trained on upright images, **(D-F)** show data for fc6 of networks trained on images rotated 22.5° counter-clockwise of upright, **(G-I)** show data for fc6 of networks trained on images rotated 45° counter-clockwise of upright. For each group of networks, panels **(A,D,G)** show FI plotted as a function of orientation, with error bars reflecting standard deviation across four networks with the same training image set **(B,E,H)** show distribution of fc6 unit tuning centers, combining data across networks **(C,F,I)** show concentration parameter (k) versus center for individual units.

The distribution of single-unit tuning properties also shifted with training set statistics. In the upright-trained model, the highest proportion of units had their tuning near the cardinals, while the networks trained on 22.5° and 45° rotated images had more units with tuning at either 22.5° or 45° counter-clockwise relative to the cardinal orientations, respectively (Figure 7B,7E,7H). Additionally, for all models, the most narrowly-tuned units tended to be those that were tuned to the orientations most common in the training set (Figure 7C,7F,7I). As described above, the high number of narrowly-tuned units with their centers close to these most common orientations may underly the double-peaked shape seen in FI.

Calculating the FIB for each of these models further demonstrated how these effects emerged across the processing hierarchy. Like the pre-trained model, the models trained on upright images showed high values of the FIB-0 at middle and deep layers: models showed significantly higher FIB-0 than the randomly initialized models for pool1, conv3_1, and all layers deeper than conv3_1 (one-tailed nonparametric t-test, FDR corrected q=0.01) (Figure 8A). In contrast, the models trained on images rotated by 22.5° and 45° showed higher values for the FIB-22 and FIB-45, respectively (Figure 8B,8C). In models trained on images rotated by 22.5°, the FIB-22 significantly exceeded that of the random models at pool2 and all layers deeper than pool2, with the exception of conv3_3 (one-tailed nonparametric t-test, FDR corrected q=0.01). For the models trained on 45° rotated images, the FIB-45 significantly exceeded that of the random models for conv3_1 and all layers deeper than conv3_1 (one-tailed nonparametric t-test, FDR corrected q=0.01).

**Figure 8.**
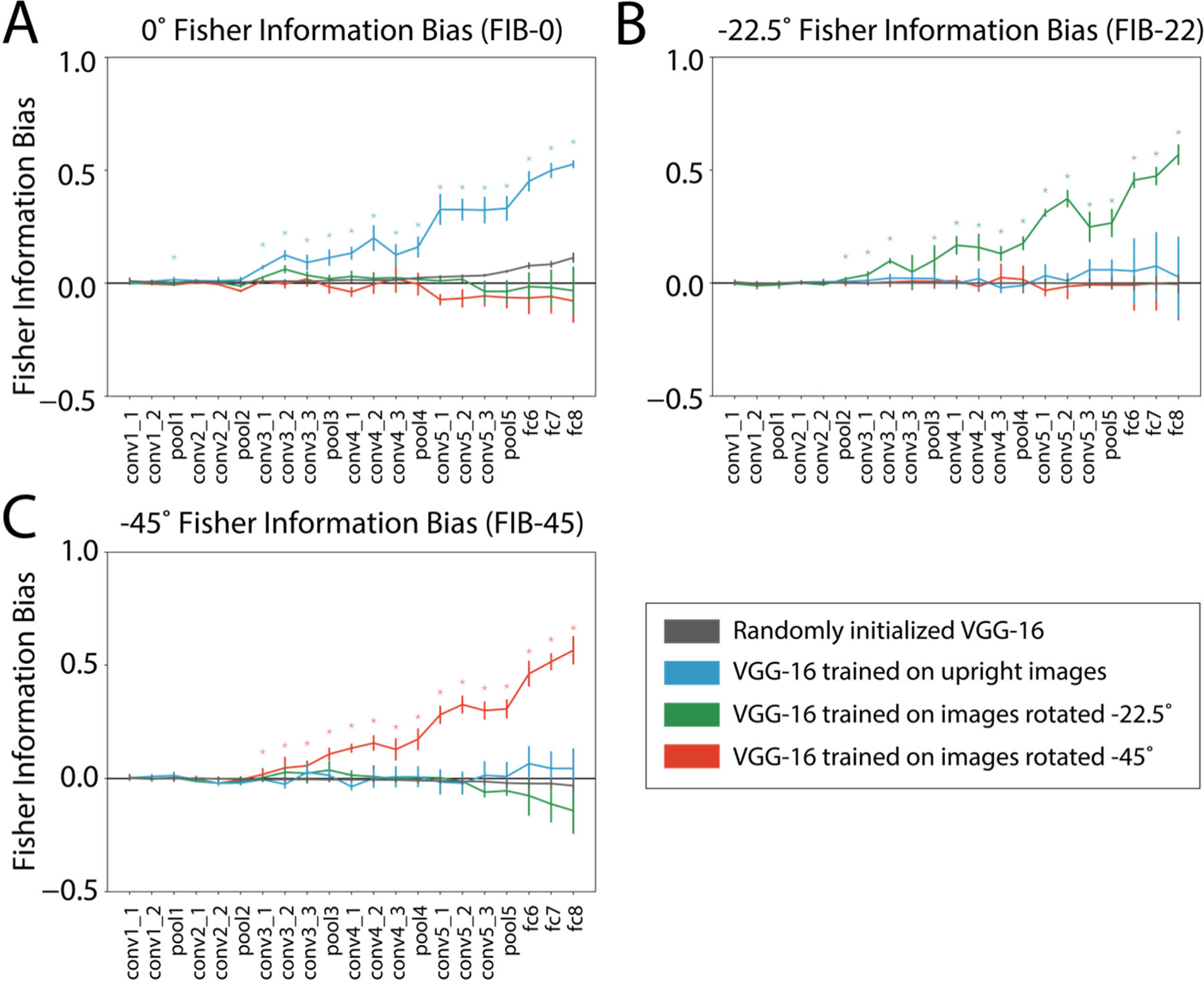
Networks shows biases in orientation discriminability that are consistent with training set statistics. FIB-0, FIB-22, and FIB-45 represent the relative value of FI at cardinal orientations, 22.5° counter-clockwise of cardinals, and 45° counter-clockwise of cardinals, respectively, relative to a baseline (see *Methods, Fisher information bias)*. Panels show **(A)** FIB-0, **(B)** FIB-22, and **(C)** FIB-45 for models trained on each rotated version of ImageNet (colored), and randomly initialized models (gray). Colored asterisks indicate layers for which the models corresponding to that color had significantly higher FIB than the random models (one-tailed nonparametric t-test, FDR corrected q=0.01). Error bars represent the standard deviation of the FIB over four initializations of each model and four evaluation image sets.

### Emergence of biases during training

Though we have demonstrated that biases were measurable in each fully trained model, this leaves open the question of whether and how these biases relate to the model’s performance at its primary task (i.e. categorizing object images). One way to address this is by examining how biases emerged over time during the training process. We hypothesized that if biases contribute to the model’s ability to learn the object classification task, then we should observe strong biases early in training, before task performance has plateaued. On the other hand, if cardinal biases are only detectable after the model has reached asymptotic performance, this would argue against the idea that they critically contribute to task performance. As shown in Figure 9, our results support the first possibility. For a network trained on upright images, Fisher information profiles measured as early as 50,000 steps into the training process already demonstrated pronounced peaks near the cardinal orientations (Figure 9A; for reference, our main analyses were performed at 400,000 steps). These peaks were not present in the model prior to any training (compare to Figure 2A, gray lines). Interestingly, these peaks in FI were larger in magnitude and shifted further away from the cardinal orientations than those measured at later timesteps (as far as 15° shifted). To investigate why this was the case, we also analyzed the tuning properties of single units at several steps during training. This revealed that, as in the fully-trained model, network layers analyzed early in training had many narrowly-tuned (high k) units with tuning centers at the cardinal orientations (Figure 9B). However, in addition, there were many units with narrow tuning whose centers lay a few degrees off of the cardinal orientations (additional “peaks” in the scatter plot in Figure 9B). The presence of these additional, off-cardinal-preferring, units is likely related to the shape of the FI measured at early timesteps. We speculate on this issue further in the *Discussion*.

**Figure 9.**
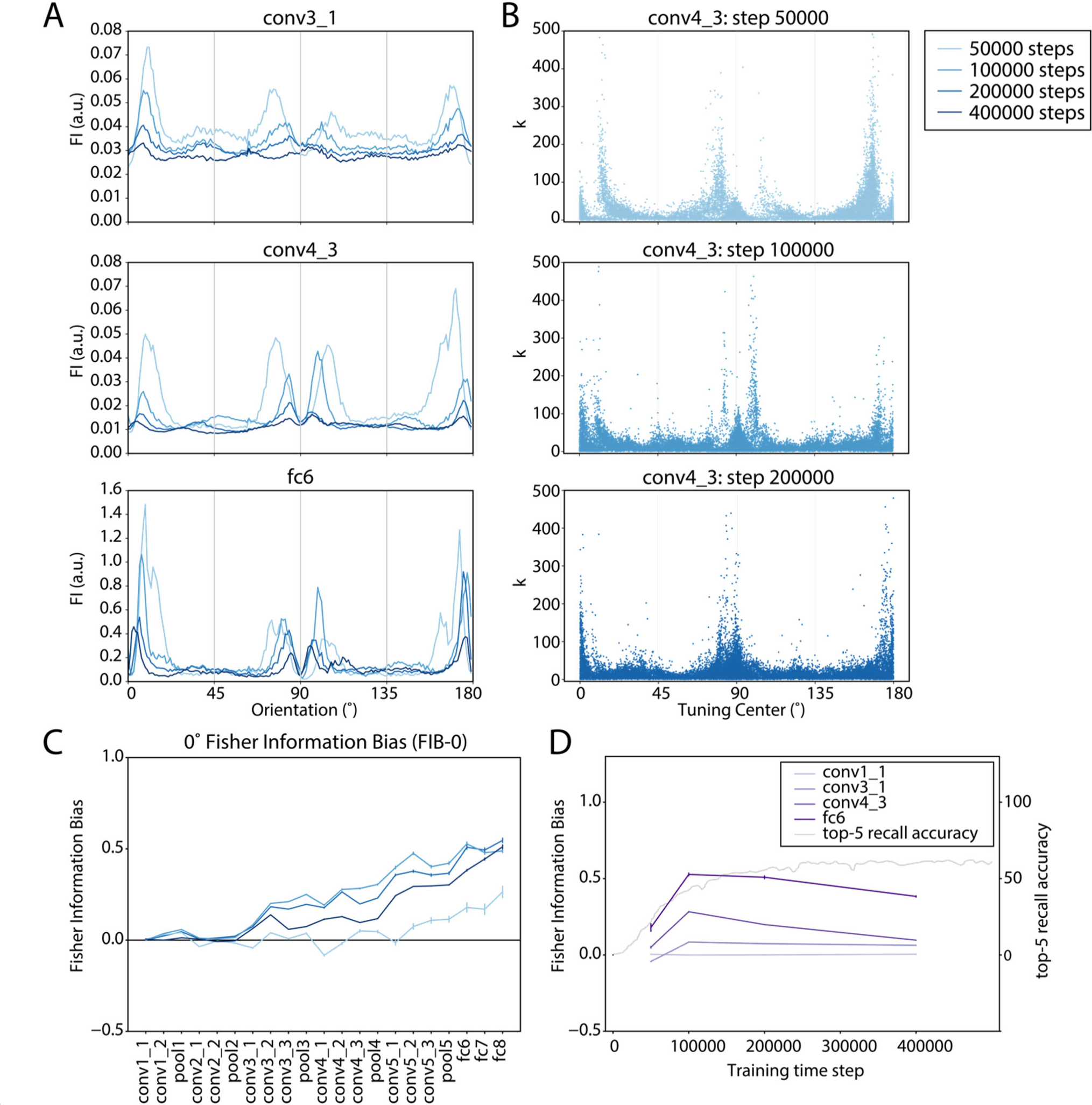
Biases in Fisher information and unit tuning properties over the course of training on upright images. **(A)** Fisher information for three example layers, at several timepoints during training (shades of blue). For comparison, analyses in Figures 2 and 3 were performed at step 400,000 (darkest blue line). **(B)** Concentration parameter (k) versus tuning center, for individual units at conv4_3, plotted for several timepoints during training. Data have been down-sampled to a maximum of 20,000 points for visualization purposes. **(C)** FIB-0 across layers plotted for several timepoints (shades of blue). Error bars reflect ±1 standard deviation of the measure across 4 evaluation image sets. **(D).** FIB-0 is plotted as a function of time, for several example layers (purple lines). Light gray line indicates model performance (top-5 recall accuracy), after smoothing with a Gaussian kernel. Error bars reflect ±1 standard deviation of the measure across 4 evaluation image sets.

As a quantitative measure of how cardinal bias evolved over time, we computed the FIB-0 as described in previous sections (Figure 9C, 9D). At all timesteps, FIB-0 tended to increase with depth in the network, as in the fully-trained model. Comparing values across training time, FIB-0 was generally maximal at 100,000 steps into the training process. We note, however, that this measure was computed by averaging the FI within 20° bins near the cardinals (see *Methods, Fisher information bias)*, and the very shifted peaks we saw at 50,000 steps were outside this window. Thus, this measure does not capture the high degree of non-uniformity we see at 50,000 steps. With this in mind, the overall amount of bias generally appeared to be highest early in training, before performance began to plateau, and decreased with additional training. At the same time, more training resulted in peaks in FI that were closer to the exact cardinal orientations, reflecting a better match to the prior distribution of orientations in the training set. It is possible that training this model for longer durations or modifying its hyperparameters to reach better plateau performance would result in further shifts in the FI peaks. However, we note that the pre-trained VGG-16 model discussed earlier (Silberman & Guadarrama, 2016) showed a similarly shaped FI curve that had peaks around the same orientations as the latest timestep of our model. Thus, it is not likely that the shape we observed is solely dependent on our choice of hyperparameters or training duration.

## Discussion

We investigated whether CNNs trained to perform object classification exhibit biased orientation representations that reflect non-uniformities in the statistics of the training set images. We found that middle and deep layers of a pre-trained VGG-16 network (Silberman & Guadarrama, 2016; Simonyan & Zisserman, 2014) represented orientation with higher discriminability near the cardinal (vertical and horizontal) orientations, with relatively lower discriminability around oblique (diagonal) orientations. Bias was also seen in the tuning properties of single units in the network: there was an over-representation of units that preferred the cardinal orientations, and units tuned to the cardinal orientations had narrower tuning profiles. Furthermore, when we trained models with the same architecture on rotated versions of ImageNet, each of these effects shifted by a predictable amount, such that discriminability was highest near whichever orientations were most common in the network’s training set. These results demonstrate that general visual experience with non-uniform image statistics is sufficient to produce, in a neural network, the same kind of biases that are observed for low-level feature representations in a range of animal species.

In general, the strength of the biases we measured tended to increase with depth in each network, showing little or no bias in the first 4-6 layers (Figure 3, Figure 4B, Figure 8). In primates, neural correlates of the oblique effect, reflected by an over-representation of cardinal-tuned neurons, have been shown in V1 (Celebrini, Thorpe, Trotter, & Imbert, 1993; De Valois, William Yund, & Hepler, 1982; Mansfield, 1974), V2 (Shen et al., 2014), and IT cortex (Vogels & Orban, 1994). To relate these physiology findings to our results, we can consider a recent finding that for a similar network, VGG-19, the ability of network activations to explain data from primate V1 was best at an intermediate layer, conv3_1, suggesting that earlier layers of the model might be more analogous to processing in the retina and/or lateral geniculate nucleus (Cadena et al., 2019). Therefore, our observation that bias did not emerge until the middle layers of the VGG-16 model is roughly consistent with a cortical origin for the oblique effect. Beyond the middle layers, we found that Fisher information bias (Figure 3, Figure 4B), continued to increase with depth in the network. At the same time, we note that another metric of orientation separability, in which we computed a t-statistic of distances in principal component space (Figure S2), showed similar values of bias at middle and late model layers. Thus, the extent to which biases are magnified versus merely reproduced at later model layers is not entirely clear. In either case, all of the methods that we employed suggest that late layers of the model consistently show strong cardinal biases. This finding is consistent with some behavioral and physiological results suggesting that the primate oblique effect may be dependent on higher-order processing beyond V1 (Shen et al., 2014; Westheimer, 2003).

Another property of the biases we observed was that the FI measured in deep layers of each network tended to peak just a few degrees off of the orientations that were most common in the training set, with a dip at the precise locations of the most common orientations (Figure 2A). As discussed in the *Results*, this double-peaked shape follows from the fact that FI is highest on the flanks of tuning curves, and many narrowly-tuned units in deep layers tended to have their centers around the most common orientations. However, this finding is not generally reflected in human psychophysics, in which the ability to make small orientation discriminations tends to show a single maximum around each of the cardinal orientations (Appelle, 1972; Girshick et al., 2011). One potential reason for this apparent discrepancy is that in this experiment, we were able to present a relatively large number of images (8640 per image set) to the CNN, with images finely spaced by 1° steps in orientation, whereas psychophysics experiments typically present fewer images at more coarsely spaced orientations (Caelli, Brettel, Rentschler, & Hilz, 1983; Girshick et al., 2011; Westheimer, 2003). Additionally, we were measuring directly from every unit without any additional sources of downstream noise or interference, which may have made the double-peaked shape of Fisher information more apparent than it would be when estimating orientation thresholds from behavior (Butts & Goldman, 2006). It is also possible that this qualitative difference between the FI curves we measured and the shape of human discriminability functions represents an actual difference between visual processing in CNNs and primates. More extensive behavioral experiments may be needed to resolve this.

From an efficient coding perspective, this double-peaked shape represents a divergence of FI from the shape predicted based on mutual information maximization (Figure 1D; Ganguli & Simoncelli, 2011; Wei & Stocker, 2015). This implies that the network is not optimizing based on mutual information but is optimizing another criterion. One interpretation is that rather than being optimized to precisely represent the orientation of stimuli, the network is optimized for simply detecting the most common stimuli, without as much regard to their precise orientation. On this account, the large number of units with their preferred orientations close to the cardinals would serve as effective detectors of cardinally-oriented stimuli but contribute less to the discrimination of these stimuli (Regan & Beverley, 1985). This interpretation raises the possibility that in the context of object categorization, detecting items with a commonly-encountered orientation is more important than discriminating their orientation.

To further investigate the functional role of these biases, we examined the time at which biases emerged during training. In a model we trained on upright images, cardinal biases were detectable early in training, before performance had begun to plateau (Figure 9). This result is consistent with the idea that the biases may contribute to improvements in task performance, rather than emerging only as a late stage byproduct of high performance. Another implication of this result is that the learned features that allow the model to achieve its maximal task performance may not be strictly required for orientation biases to emerge. Rather, it may be the case that as soon as the network begins to represent a feature such as orientation, the resulting representations exhibit bias. Indeed, the overall measure of cardinal information bias declined from step 100,000 to step 400,000 (Figure 9D), suggesting that biases are initially quite pronounced and become scaled back over the course of training.

At the same time, the double-peaked shape in FI was even more pronounced early in training, with the peaks further offset from the cardinal axes (Figure 9A). As described earlier in this section, this double-peaked shape may be non-optimal in terms of efficient coding. Thus, it may be the case that training has the dual effects of reducing the magnitude of bias and making the network’s representations more closely approximate an efficient encoding of the prior distribution. This process apparently unfolds in parallel to the model’s improvement in task performance, leaving open the question of what relationship between these processes is. One interpretation is that optimal orientation coding directly impacts the model’s learning process, contributing to its ability to accurately classify objects. Another interpretation is that both the changes in orientation bias and the improvement in object categorization performance are reflections of the same underlying process, in which the network is developing a more accurate, complete representation of the training set distribution. Further experiments will be needed to conclusively resolve this.

Our findings may also provide some indication of which architectural properties of the VGG-16 network are required for cardinal biases to emerge. First, we found that biases tended to emerge around the 5^th^ layer of the model and increase with depth in the network (Figure 3).

The fact that there is a minimum depth requirement for biases to emerge may be related to the small convolutional kernels in the VGG-16 model (3x3 pixels), potentially suggesting that a large spatial receptive field is required for the emergence of orientation biases. A potential extension of this work would be to compare our results with another network architecture having a larger convolutional filter size, and examine whether biases are present at a shallower depth in such a model (e.g. AlexNet; Krizhevsky, Sutskever, & Hinton, 2012). The observation that biases further increase with depth in the model may further suggest that repeated convolutions, spatial pooling operations and fully connected layers result in amplification of the biases. Additionally, more general properties of the convolutional architecture used in VGG-16 might be required for the emergence of the oblique effect. For example, the sharing of weights across units with different spatial selectivity may be a critical feature that enables learning of orientation representations that efficiently encode the training set statistics. The use of a nonlinear activation function may also be key. Manipulating these core aspects of the model is beyond the scope of the present paper, but in future work it would be useful to explore biases in visual representations across a wider range of network architectures. For instance, if it were found that the use of a convolutional architecture, roughly analogous to the architecture of the primate visual system, is critical for cardinal biases to emerge, this might provide insight into the functional relevance of this property in the primate brain.

Finally, we also observed weak evidence for a cardinal bias in FI measured from the deep layers of a random network with no training (Figure 3, Figure 8A). This may indicate that some aspect of the model’s architecture, such as its use of a square image grid, square convolutional kernels, and pooling operations over square image regions, introduced an intrinsic cardinal reference frame. However, the possible presence of such a reference frame cannot account for the effects we observed for several reasons. First, the magnitude of the FIB-0 was 5x lower for the deepest layer of the random models as compared to the trained-upright models, and the random models did not show an over-representation of cardinal-tuned units, while the upright-trained models did (Figure 2B, Figure 7B, Figure S1). This suggests that the network response properties underlying any intrinsic cardinal FI bias were different than those underlying the experience-driven biases we observed. Second, the magnitude of the shifted biases we measured in models trained on rotated images were of similar magnitude to the cardinal biases we measured in models trained on upright images (Figure 8), which demonstrates that having an intrinsic reference frame that matches the orientation distribution of training images is not required for a substantial bias to emerge. These results suggest that training may be able to override some intrinsic response properties of CNNs. However, they also highlight the general importance of examining the biases inherent in CNNs before making analogies to the visual system.

These findings also have general relevance for the use of CNNs in vision research. First, our results show that a popular CNN model exhibits a form of the classical oblique effect, suggesting that this key aspect of low-level primate vision is reproduced by the model. This adds to a growing body of work demonstrating similarities between deep neural networks and the brains and behavior of primates (Kubilius, Bracci, & Op de Beeck, 2016; Pospisil, Pasupathy, & Bair, 2018; Rideaux & Welchman, 2020; Ward, 2019; Yamins et al., 2014). Second, we have demonstrated that non-uniformities in the statistics of training set images can dramatically influence the feature representations that are learned by a CNN. Specifically, image features that are over-represented during training are likely to be more discriminable by the trained network, which may lead to a performance advantage for processing certain stimuli over others. Accounting for such influences is critical for avoiding unwanted algorithmic biases, particularly in modeling high-level visual functions such as face recognition (Buolamwini & Gebru, 2018; Cavazos, Phillips, Castillo, & O’Toole, 2019; Klare, Burge, Klontz, Vorder Bruegge, & Jain, 2012).

Overall, our results suggest that the classical oblique effect is reproduced in a CNN trained to perform object recognition on an image set containing an over-representation of cardinal orientations. Furthermore, a rotated version of this bias can be induced by training a CNN on rotated versions of these same images. These results indicate that general visual experience, without the presence of an innate bias that matches the viewed orientation distribution, is sufficient to induce the formation of orientation biases, providing support for an experience-driven account of the oblique effect.

## Acknowledgements

Funded by NEI R01-EY025872 to JTS, and NIMH Training Grant in Cognitive Neuroscience (T32-MH020002) to MMH. We thank the San Diego Supercomputer Center and UCSD’s Social Sciences uting Facility for computing support, and the NVIDIA corporation for donation of a Quadro GPU that was used in this research.

## Supplementary Figures

**Figure S1.**
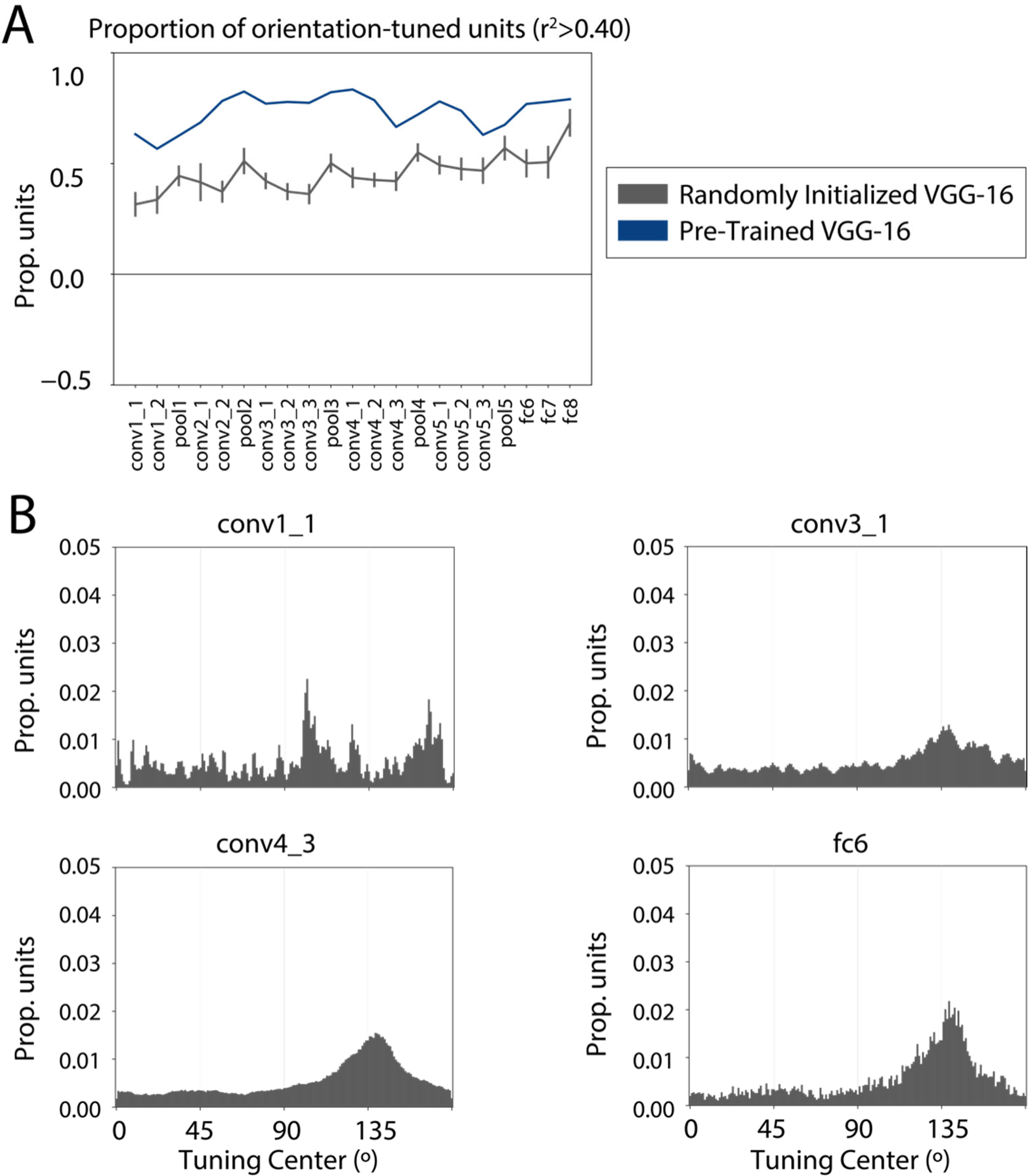
**(A)** Proportion of units in each layer that were well-fit by a Von Mises function (see *Methods, Single-unit tuning analysis*), for a pre-trained VGG-16 model (navy blue) and randomly initialized models with no training (gray). Error bars on the gray line reflect standard deviation across four different random initializations of the model. **(B)** Distribution of unit tuning centers for the randomly initialized models (distributions are combined across four different random initializations of the model).

**Figure S2.**
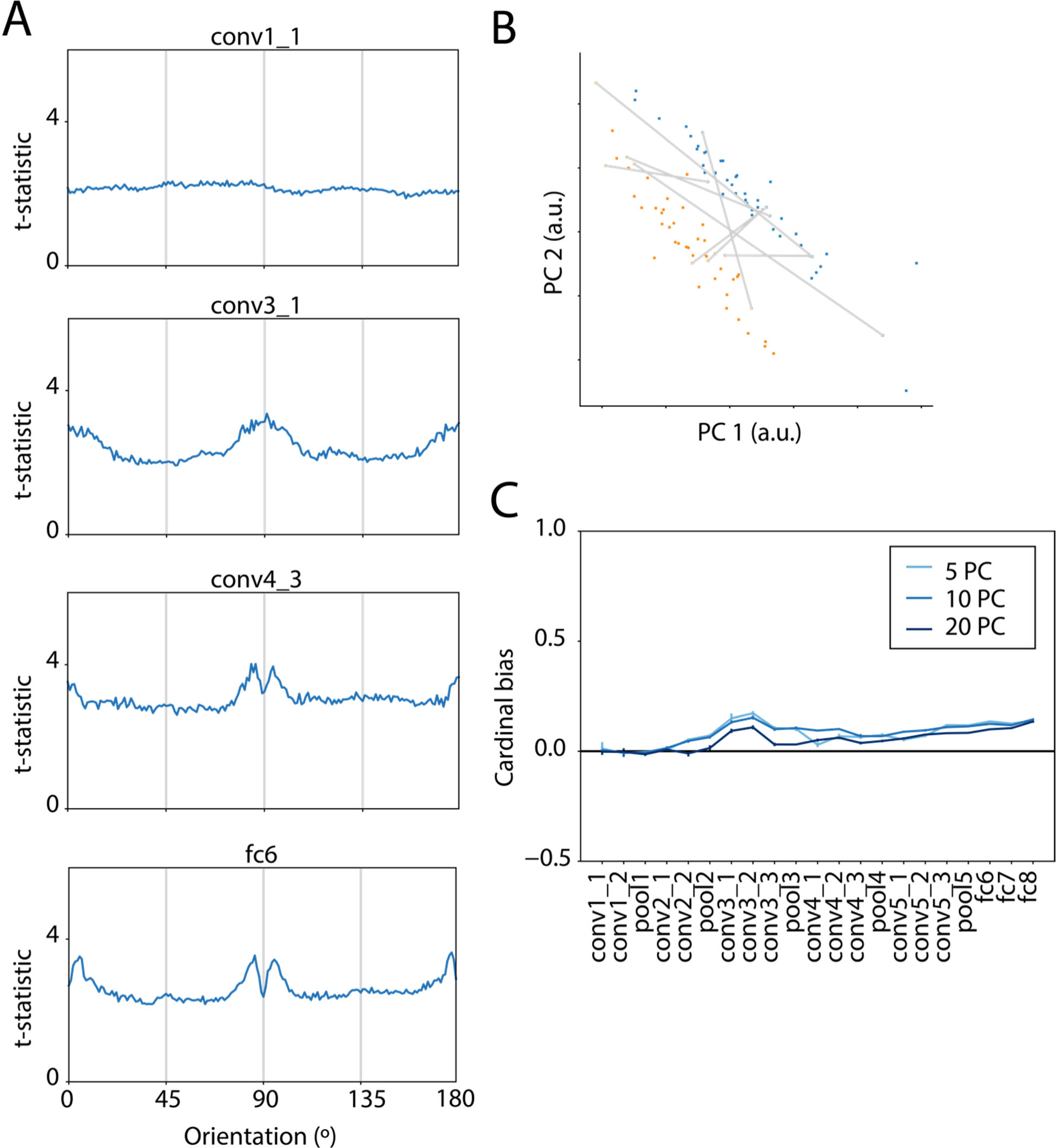
**(A)** T-statistic for distances between points corresponding to different orientations, in principal component space. Calculation was done using the first 10 PCs (see *Methods, Multivariate analyses)*. **(B)** Schematic of how the t-statistic distance measure was calculated in PC space: orange and blue dots indicate representations of two nearby orientations, gray lines indicate distances between points corresponding to different orientations (note that there are 2304 such pairwise distances in total; not all distances are drawn). The t-statistic corresponds to the mean of these distances divided by the standard deviation of the distances. **(C)** A measure equivalent to FIB (see *Methods, Fisher information bias)* for the t-statistic distance is plotted for each layer, comparing results with 5, 10 or 20 principal components. Error bars reflect ±1 standard deviation of the measure across 4 evaluation image sets.

## References

Abadi, M., Barham, P., Chen, J., Chen, Z., Davis, A., Dean, J., … Zheng, X. (2016). TensorFlow: A system for large-scale machine learning. ArXiv. Retrieved from http://arxiv.org/abs/1605.08695

Abbott, L. F., & Dayan, P. (1999). The effect of correlated variability on the accuracy of a population code. In Neural Computation (Vol. 11). https://doi.org/10.1162/089976699300016827

Appelle, S. (1972). Perception and discrimination as a function of stimulus orientation: The “oblique effect” in man and animals. Psychological Bulletin, 78(4), 266–278. https://doi.org/10.1037/h0033117

Barlow, H. B. (1961). Possible Principles Underlying the Transformations of Sensory Messages. In Sensory Communication (pp. 217–234). https://doi.org/10.7551/mitpress/9780262518420.003.0013

Bauer, J. A., Owens, D. A., Thomas, J., & Held, R. (1979). Monkeys Show an Oblique Effect. Perception, 8(3), 247–253. https://doi.org/10.1068/p080247

Benjamini, Y., & Yekutieli, D. (2001). The control of the false discovery rate in multiple testing under dependency. Annals of Statistics, Vol. 29, pp. 1165–1188. https://doi.org/10.1214/aos/1013699998

Blakemore, C., & Cooper, G. F. (1970). Development of the brain depends on the visual environment. Nature, 228(5270), 477–478. https://doi.org/10.1038/228477a0

Buolamwini, J., & Gebru, T. (2018). Gender Shades: Intersectional Accuracy Disparities in Commercial Gender Classification *. In Proceedings of Machine Learning Research (Vol. 81). Retrieved from PMLR website: http://proceedings.mlr.press/v81/buolamwini18a.html

Butts, D. A., & Goldman, M. S. (2006). Tuning Curves, Neuronal Variability, and Sensory Coding. PLoS Biology, 4(4), e92. https://doi.org/10.1371/journal.pbio.0040092

Cadena, S. A., Denfield, G. H., Walker, E. Y., Gatys, L. A., Tolias, A. S., Bethge, M., & Ecker, A. S. (2019). Deep convolutional models improve predictions of macaque V1 responses to natural images. PLoS Computational Biology, 15(4), e1006897. https://doi.org/10.1371/journal.pcbi.1006897

Caelli, T., Brettel, H., Rentschler, I., & Hilz, R. (1983). Discrimination thresholds in the two-dimensional spatial frequency domain. Vision Research, 23(2), 129–133. https://doi.org/10.1016/0042-6989(83)90135-9

Cavazos, J. G., Phillips, P. J., Castillo, C. D., & O’Toole, A. J. (2019). Accuracy comparison across face recognition algorithms: Where are we on measuring race bias? Retrieved from http://arxiv.org/abs/1912.07398

Celebrini, S., Thorpe, S., Trotter, Y., & Imbert, M. (1993). Dynamics of orientation coding in area VI of the awake primate. Visual Neuroscience, 10(5), 811–825. https://doi.org/10.1017/S0952523800006052

Cichy, R. M., & Kaiser, D. (2019). Deep Neural Networks as Scientific Models. Trends in Cognitive Sciences, 23(4), 305–317. https://doi.org/10.1016/j.tics.2019.01.009

Coppola, D. M., Purves, H. R., McCoy, A. N., & Purves, D. (1998). The distribution of oriented contours in the real world. Proceedings of the National Academy of Sciences of the United States of America, 95(7), 4002–4006. https://doi.org/10.1073/pnas.95.7.4002

Coppola, D. M., & White, L. E. (2004). Visual experience promotes the isotropic representation of orientation preference. Visual Neuroscience, 21(1), 39–51. https://doi.org/10.1017/s0952523804041045

De Valois, R. L., William Yund, E., & Hepler, N. (1982). The orientation and direction selectivity of cells in macaque visual cortex. Vision Research, 22(5), 531–544. https://doi.org/10.1016/0042-6989(82)90112-2

Deng, J., Dong, W., Socher, R., Li, L.-J., Kai Li, & Li Fei-Fei. (2009). ImageNet: A large-scale hierarchical image database. In 2009 IEEE Conference on Computer Vision and Pattern Recognition. https://doi.org/10.1109/CVPRW.2009.5206848

Ganguli, D., & Simoncelli, E. P. (2011). Implicit encoding of prior probabilities in optimal neural population (Vol. 23). Retrieved from MIT Press website: http://www.nips.cc

Girshick, A. R., Landy, M. S., & Simoncelli, E. P. (2011). Cardinal rules: visual orientation perception reflects knowledge of environmental statistics. Nature Neuroscience, 14(7), 926–932. https://doi.org/10.1038/nn.2831

Hirsch, H. V. B., & Spinelli, D. N. (1970). Visual experience modifies distribution of horizontally and vertically oriented receptive fields in cats. Science, 168(3933), 869–871. https://doi.org/10.1126/science.168.3933.869

Hoy, J. L., & Niell, C. M. (2015). Layer-specific refinement of visual cortex function after eye opening in the awake mouse. The Journal of Neuroscience : The Official Journal of the Society for Neuroscience, 35(8), 3370–3383. https://doi.org/10.1523/JNEUROSCI.3174-14.2015

Jain, A. K., & Farrokhnia, F. (1991). Unsupervised texture segmentation using Gabor filters. Pattern Recognition, 24(12), 1167–1186. https://doi.org/10.1016/0031-3203(91)90143S

Kell, A. J., & McDermott, J. H. (2019, April 1). Deep neural network models of sensory systems: windows onto the role of task constraints. Current Opinion in Neurobiology, Vol. 55, pp. 121–132. https://doi.org/10.1016/j.conb.2019.02.003

Klare, B. F., Burge, M. J., Klontz, J. C., Vorder Bruegge, R. W., & Jain, A. K. (2012). Face recognition performance: Role of demographic information. IEEE Transactions on Information Forensics and Security, 7(6), 1789–1801. https://doi.org/10.1109/TIFS.2012.2214212

Kreile, A. K., Bonhoeffer, T., & Hübener, M. (2011). Altered visual experience induces instructive changes of orientation preference in mouse visual cortex. Journal of Neuroscience, 31(39), 13911–13920. https://doi.org/10.1523/JNEUROSCI.2143-11.2011

Krizhevsky, A., Sutskever, I., & Hinton, G. E. (2012). ImageNet Classification with Deep Convolutional Neural Networks. In Advances in Neural Information Processing Systems (Vol. 25). Retrieved from http://code.google.com/p/cuda-convnet/

Kubilius, J., Bracci, S., & Op de Beeck, H. P. (2016). Deep Neural Networks as a Computational Model for Human Shape Sensitivity. PLOS Computational Biology, 12(4), e1004896. https://doi.org/10.1371/journal.pcbi.1004896

Leventhal, A. G., & Hirsch, H. V. B. (1975). Cortical effect of early selective exposure to diagonal lines. Science, 190(4217), 902–904. https://doi.org/10.1126/science.1188371

Leventhal, A. G., & Hirsch, H. V. B. (1980). Receptive-field properties of different classes of neurons in visual cortex of normal and dark-reared cats. Journal of Neurophysiology, 43(4), 1111–1132. https://doi.org/10.1152/jn.1980.43.4.1111

Li, B., Peterson, M. R., & Freeman, R. D. (2003). Oblique Effect: A Neural Basis in the Visual Cortex. Journal of Neurophysiology, 90(1), 204–217. https://doi.org/10.1152/jn.00954.2002

Mansfield, R. J. W. (1974). Neural basis of orientation perception in primate vision. Science, 186(4169), 1133–1135. https://doi.org/10.1126/science.186.4169.1133

Pospisil, D. A., Pasupathy, A., & Bair, W. (2018). ’Artiphysiology’ reveals V4-like shape tuning in a deep network trained for image classification. ELife, 7. https://doi.org/10.7554/eLife.38242

Regan, D., & Beverley, K. I. (1985). Postadaptation orientation discrimination. Journal of the Optical Society of America. A, Optics and Image Science, 2(2), 147–155. Retrieved from http://www.ncbi.nlm.nih.gov/pubmed/3973752

Rideaux, R., & Welchman, A. E. (2020). But still it moves: Static image statistics underlie how we see motion. Journal of Neuroscience, 40(12), 2538–2552. https://doi.org/10.1523/JNEUROSCI.2760-19.2020

Russakovsky, O., Deng, J., Su, H., Krause, J., Satheesh, S., Ma, S., … Fei-Fei, L. (2015). ImageNet Large Scale Visual Recognition Challenge. International Journal of Computer Vision, 115(3), 211–252. https://doi.org/10.1007/s11263-015-0816-y

Shen, G., Tao, X., Zhang, B., Smith, E. L., Chino, Y. M., & Chino, Y. M. (2014). Oblique effect in visual area 2 of macaque monkeys. Journal of Vision, 14(2). https://doi.org/10.1167/14.2.3

Silberman, N., & Guadarrama, S. (2016). TensorFlow-Slim image classification model library. Simonyan, K., & Zisserman, A. (2014). Very Deep Convolutional Networks for Large-Scale Image Recognition. Retrieved from http://arxiv.org/abs/1409.1556

Vogels, R., & Orban, G. A. (1994). Activity of inferior temporal neurons during orientation discrimination with successively presented gratings. Journal of Neurophysiology, 71(4), 1428–1451. https://doi.org/10.1152/jn.1994.71.4.1428

Wainwright, M. J. (1999). Visual adaptation as optimal information transmission. Vision Research, 39(23), 3960–3974. https://doi.org/10.1016/S0042-6989(99)00101-7

Ward, E. J. (2019). Exploring perceptual illusions in deep neural networks. Journal of Vision, 19(10), 34b. https://doi.org/10.1167/19.10.34b

Wei, X.-X., & Stocker, A. A. (2015). A Bayesian observer model constrained by efficient coding can explain “anti-Bayesian” percepts. Nature Neuroscience, 18(10), 1509–1517. https://doi.org/10.1038/nn.4105

Westheimer, G. (2003). Meridional anisotropy in visual processing: Implications for the neural site of the oblique effect. Vision Research, 43(22), 2281–2289. https://doi.org/10.1016/S0042-6989(03)00360-2

Yamins, D., Hong, H., Cadieu, C. F., Solomon, E. A., Seibert, D., & DiCarlo, J. J. (2014). Performance-optimized hierarchical models predict neural responses in higher visual cortex. Proceedings of the National Academy of Sciences, 111(23), 8619–8624. https://doi.org/10.1073/pnas.1403112111

